# Glassiness in cellular Potts model of biological tissue is controlled by disordered energy landscape

**DOI:** 10.1101/2020.08.27.270488

**Authors:** Souvik Sadhukhan, Saroj Kumar Nandi

## Abstract

Glassy dynamics in a confluent monolayer is indispensable in morphogenesis, wound healing, bronchial asthma, and many others; a detailed theoretical understanding for such a system is, therefore, important. We combine numerical simulations of a cellular Potts model and an analytical study based on random first order transition (RFOT) theory of glass, develop a comprehensive theoretical framework for a confluent glassy system, and show that glassiness is controlled by the underlying disordered energy landscape. Our study elucidates the crucial role of geometric constraints in bringing about two distinct regimes in the dynamics, as the target perimeter *P*_0_ is varied. The extended RFOT theory provides a number of testable predictions that we verify in our simulations. The unusual sub-Arrhenius relaxation results from the distinctive interaction potential arising from the perimeter constraint in a regime controlled by geometric restriction. Fragility of the system decreases with increasing *P*_0_ in the low-*P*_0_ regime, whereas the dynamics is independent of *P*_0_ in the other regime. The mechanism, controlling glassiness in a confluent system, is different in our study in comparison with vertex model simulations, and can be tested in experiments.

---

Hallmarks of glassiness in collective motion in an epithelial monolayer [1–3] are important in morphogenesis [4–7], wound healing [8–11], vertibrate body axis elongation [12], tumor progression [6, 13], bronchial asthma [14, 15], and many other. Developing a detailed theoretical framework for the glassy dynamics in such systems is, therefore, an important task. Inspired by the physics of soap bubbles, vertex-based models [16–20] that represent individual cells by polygons have provided important insights into the dynamics of such systems [21–26]. Within these models, the cellular perimeter between vertices is either straight by construction or has a constant curvature, whereas in experiments it deviates arbitrarily from a straight line [19, 27, 28]. How this deviation affects the dynamics remains unknown. Moreover, the *T*1 transition or the neighbor exchange process, that is crucial for dynamics in such systems, needs to be externally imposed and sensitive to model parameters [15].

The lattice-based cellular Potts models (CPM) [29–31] define another important class of models for cellular dynamics and has been applied to single and collective cellular behavior [32–35], cell sorting [29, 30], dynamics on patterned surfaces [36], gradient sensing [36, 37] etc. The primary difference between CPM and vertex-based models lies in the details of energy minimization [38]. Two crucial aspects of CPM, however, make it advantageous over vertex-based models: it allows simulation of arbitrary cell perimeters, and *T*1 transitions are naturally included within CPM. Despite the widespread applicability of CPM, its glassy aspects remain relatively unexplored. To the best of our knowledge, there exists only one such simulation study [39], which however did not consider the perimeter constraint and, as we show below, models with and without this constraint are qualitatively different. Our focus in this work is the equilibrium system. Although biological systems are inherently out of equilibrium and activity is crucial, it is important to first understand the behavior of an equilibrium system in the absence of activity, which can be included later [40].

Simulation studies of vertex-based models have established a rigidity transition that controls the glassy dynamics and the observed shape index (average ratio of perimeter to square root of area) is the structural order parameter of glass transition [14, 23, 24]. We show that these results are *not* generic for confluent systems. Our aim in this work is twofold: we bridge the gap in numerical results through detailed Monte-Carlo (MC) based simulation study of CPM in the glassy regime, and we develop RFOT theory [41, 42] for a confluent system in glassy regime and show that glassiness in such systems is controlled by the underlying disordered configurations. The other main results of this work are as follows: (i) *A*_0_ does not affect the dynamics in a confluent system,(ii) *P*_0_ that parameterizes the interaction potential, plays the role of a control parameter, (iii) geometric restriction brings about two regimes as *P*_0_ is varied; dynamics depends on *P*_0_ in the low-*P*_0_ regime and independent of *P*_0_ in the other regime, (iv) changes in fragility as well as the unusual sub-Arrhenius relaxation in such systems are results of the perimeter constraint.

## Model

The energy function for CPM (SM Sec. S1) corresponding to a confluent monolayer [31, 38] is

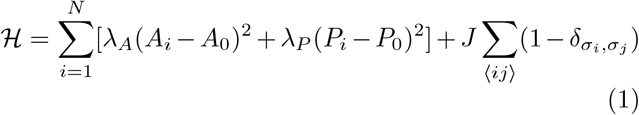

where *σ*_*i*_, (*i* ∈ 1, …, *N*) are cell indices, *N*, the total number of cells, *A*_*i*_ and *P*_*i*_ are area and perimeter of the *i*th cell, *A*_0_ and *P*_0_ are target area and target perimeter, chosen to be same for all cells. *λ*_*A*_ and *λ*_*P*_ are elastic constants related to area and perimeter constraints. 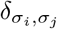 is Kronecker delta function. The first term in Eq. (1) comes from incompressibility of cells; second and third terms come from properties of cellular cortex and inter-cellular interactions [21, 43]. The third term in ℋ is proportional to *P*_*i*_ and can be included within the *λ*_*P*_ term (SM Sec. S1 S1A), however, for ease of discussion we keep it separately. CPM represents the biological processes for dynamics through an effective temperature *T*[29, 31]. We perform extensive Monte-Carlo (MC) simulations for the dynamics via an algorithm [44] that allows us to locally calculate the perimeter and makes it possible to simulate large system sizes for long times appropriate to investigate glassy dynamics. Fragmentation of cells is forbidden [45] in our simulation to minimize noise. We mainly focus on the model with *J* = 0 and get back to the model with *J* ≠ 0 and *λ*_*P*_ = 0, that was simulated in Ref. [39], later in the paper.

## I. RESULTS

### Dynamics is independent of *A*_0_

The change in energy coming from the area term alone for an MC attempt *σ*_*i*_ → *σ*_*j*_ between *i*th and *j*th cells (SM Sec. S1 S1C) is Δℋ_*area*_ = 2*λ*_*A*_(1 − *A*_*i*_ + *A*_*j*_) that is independent of *A*_0_. Since *A*_0_ dependence of dynamics can only come through Δℋ_*area*_, the dynamics becomes independent of *A*_0_. This result is a consequence of the energy function, Eq. (1), and the constraint of confluency. This argument can also be extended for a polydisperse system (SM, Sec S1 S1C). The input shape index, 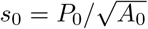, therefore cannot be a control parameter for the dynamics [14, 23, 24, 46, 47] and should be viewed as a dimensionless perimeter. *P*_0_, on the other hand, parameterizes the interaction potential and plays the role of a control parameter. Since dynamics in such a system essentially changes *A*_*i*_ and *P*_*i*_, even in a monodisperse system in steady state, we typically see a distribution for both *A*_*i*_ and *P*_*i*_ (Fig. 1(a)), implying dispersion of both cell area and interaction, leading to an effectively polydisperse system.

**FIG 1.**
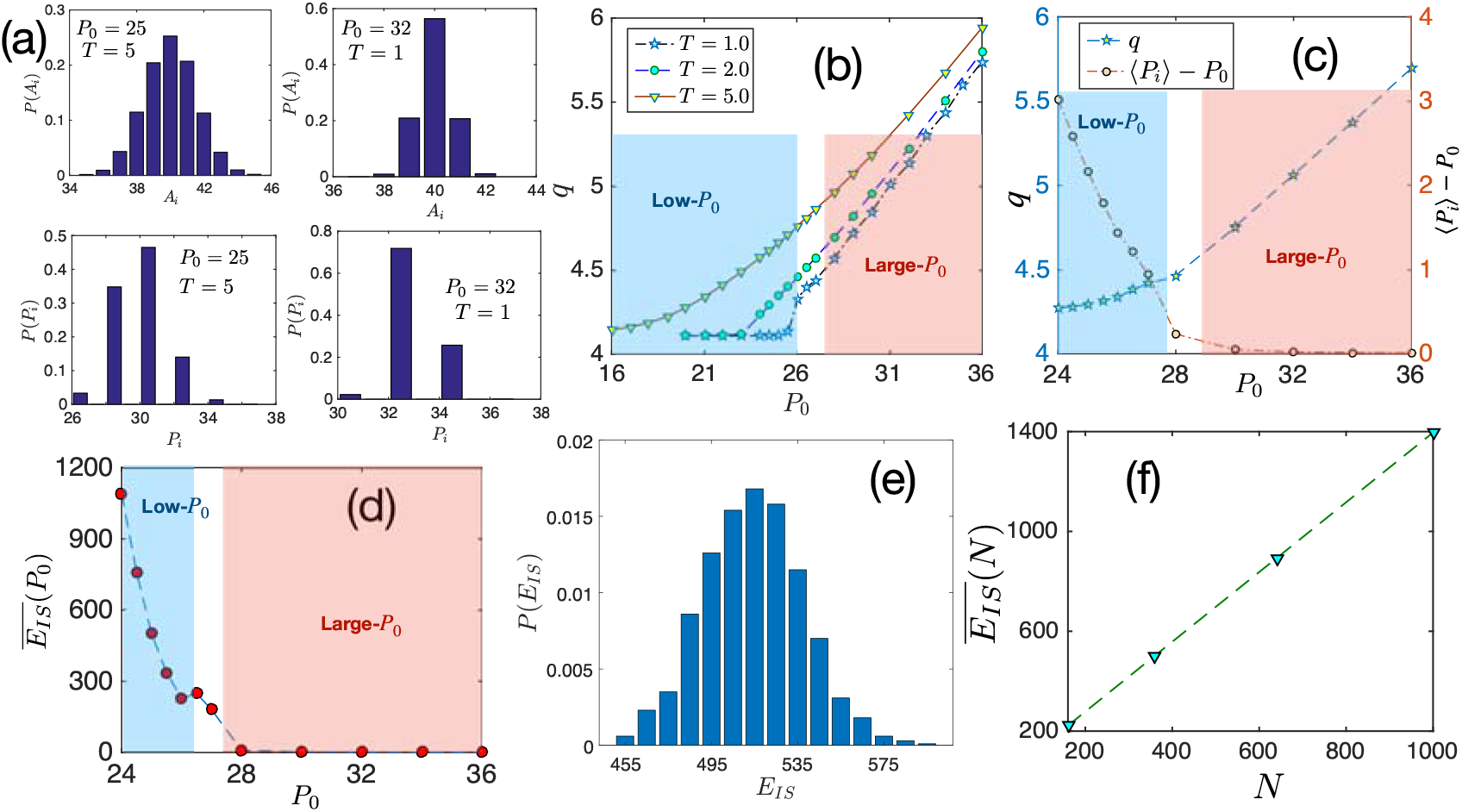
Nature of steady state and inherent structure in CPM. (a) A monodisperse system in steady state has a distribution of *A*_*i*_ and *P*_*i*_ as shown for two different *P*_0_ and *T* values quoted in the figures. This effectively leads to a polydisperse system. (b) Observed shape index, *q*, as a function of *P*_0_ at three different *T*; *P*_*min*_ = 26 in our simulations (see Materials and Methods). Lowest value of *q* is given by geometric restriction in the low-*P*_0_ regime and by *P*_0_ in the large-*P*_0_ regime. (c) *q* at *T*_*g*_ as a function of *P*_0_ tends to saturate in the low-*P*_0_ regime and increases linearly with *P*_0_ in the large-*P*_0_ regime. Right *y*-axis shows ⟨*P*_*i*_⟩ − *P*_0_ decreases linearly with increasing *P*_0_ in the low-*P*_0_ regime and then tends to zero. Each point in (b) and (c) is an average over 10^5^ *t*_0_. (d) Average inherent structure energy, 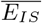, shows strong dependence on *P*_0_ in the low-*P*_0_ regime and then it becomes zero. (e) Broad distribution of *E*_*IS*_ in the low *P*_0_ regime is consistent with the existence of many local minima with varying energy. *P*_0_ = 25 and initial equilibration *T* = 4.0 for this figure. (f) Linear dependence of 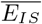 on *N* suggests extensivity of configurational entropy. Each point in (d) and (f) is an average over at least 10^3^ ensembles.

### Two different regimes of *P*_0_

The observed shape index, 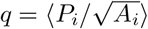, where ⟨…⟩ denotes average over all cells, tends to a constant with decreasing *P*_0_. Through simulations of vertex-based models, *q* has been interpreted as a structural order parameter of glass transition [14, 23, 24]. However, as we show below, such an interpretation is *not* generic and within CPM, the result is simply a consequence of geometric restriction. *P*_*i*_ for a fixed *A*_*i*_ has a minimum value, *P*_*min*_, that depends on geometric constraints, here confluency and underlying lattice. When *P*_0_ is below *P*_*min*_, *P*_*i*_ of most cells cannot satisfy the perimeter constraint in Eq. (1) as they remain stuck around *P*_*min*_. At high *T* fluid regime, when dynamics is fast, cell boundaries are irregular leading to larger values of *P*_*i*_ and *q*; but, at low *T* glassy regime, when dynamics is slow, cell boundaries tend to be regular leading to lower values of *P*_*i*_ and *q*. The lowest value of *q* is dictated by the geometric restriction in this low-*P*_0_ regime. Figure 1(b) shows *q* at three different *T* as a function of *P*_0_; *q* decreases with decreasing *T* at a fixed *P*_0_ and saturates to 4.11. Our interpretation is consistent with the fact that *q* in a large class of distinctly different systems has similar value [28, 48].

On the other hand, when *P*_0_ > *P*_*min*_, the large-*P*_0_ regime, most cells are able to satisfy the perimeter constraint and the lowest value of *q* is governed by *P*_0_, as deviation from *P*_0_ costs energy. We show *q* at *T*_*g*_ as a function of *P*_0_ in Fig. 1(c); *q* tends to a constant in the low-*P*_0_ regime whereas it increases linearly with *P*_0_ in the large-*P*_0_ regime. The geometric restriction is clearer in the plot of ⟨*P*_*i*_⟩ − *P*_0_; it decreases linearly with increasing *P*_0_ in the low-*P*_0_ regime and then tends to zero. The interfacial tension, defined as *γ* = ∂ℋ /∂*P*_*i*_ ∝ (*P*_*i*_ − *P*_0_) [33], is non-zero along cell boundaries in the low-*P*_0_ regime and becomes zero as *P*_0_ increases.

Effect of geometric constraint is evident in the properties of inherent structure, whose energy, *E*_*IS*_, (SM, Sec. S2) characterizes the energy landscape that are quantified through the configurational entropy, *s*_*c*_, although the exact relation among them remains unknown [49]. Since *A*_0_ does not affect dynamics, we set it to the average area. *E*_*IS*_ is non-zero in the low-*P*_0_ regime and zero in the large-*P*_0_ regime, where cells are able to satisfy the constraints in Eq. (1). 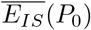, the ensemble averaged *E*_*IS*_, strongly depends on *P*_0_ in the low-*P*_0_ regime and then becomes zero (Fig. 1d). In the low-*P*_0_ regime, a broad distribution of *E*_*IS*_ [Fig. 1(e)] is consistent with the existence of many local minima with varying energy and a linear variation of 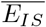 as a function of *N* (Fig. 1(f)) shows 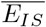, and hence *s*_*c*_, are extensive. Although *E*_*IS*_ is zero in the large-*P*_0_ regime, the inherent structure is disordered (SM Fig. S6); there exist a large number of minima with same energy separated by varying barriers. These results suggest the existence of glassy dynamics and the applicability of RFOT theory phenomenology in both regimes. If rigidity transition controls the dynamics, as in vertex models [23, 24], glass transition in the large-*P*_0_ regime should be absent. But, as we show below, the system *does* show glassiness even in this regime; this is consistent with RFOT phenomenology, where glassiness comes from the disordered energy landscape.

### RFOT theory for CPM

Within RFOT theory, a glassy system consists of mosaics of different states (SM, Sec. S3). The typical length scale, *ξ*, of such mosaics is determined from the balance of two competing contributions: the configurational entropy, *s*_*c*_, that facilitates escape of a mosaic to different states, reducing *ξ*, and the surface reconfiguration energy, Γ, that accounts for cost at the surface of two different states, increasing *ξ* [41, 42, 50, 51]. Minimizing the free energy cost for escaping the state of a certain region, one obtains *ξ* ∼ (Γ/*s*_*c*_)^1/(*d*−*θ*)^, where *d* is dimension and *θ*, an exponent relating surface area and length scale of a region. *P*_0_ sets the inter-cellular interaction potential within CPM, and therefore, *s*_*c*_ and Γ are functions of *P*_0_ as they both stem from this potential. Considering linear *T*-dependence of Γ [52], we write Γ = Ξ(*P*_0_)*T*. The relaxation dynamics within RFOT theory is governed by relaxation of these mosaics of typical length scale *ξ*. We now consider an Arrhenius-type argument where energy barrier for relaxation of a region of length scale *ξ* varies as ∼ *ξ*^*ψ*^. Taking the exponents as *θ* = *ψ* = *d*/2 [42, 52, 53], we obtain the relaxation time *τ* (SM Sec. S3) as

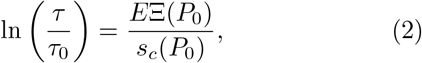

where *E* is a constant and *τ*_0_ is a high-*T* timescale that can depend on interatomic interactions and, hence, *P*_0_. Eq. (2) gives the general form of RFOT theory for the CPM; we obtain the detailed forms of Ξ(*P*_0_) and *s*_*c*_(*P*_0_) for different systems and regimes that we consider below.

### Low *P*_0_ regime

As discussed above, *P*_*i*_ for most cells are less than *P*_0_ in this regime. Figure 2(a) shows a typical configuration of cells and their centers of mass for a system, close to glass transition. The mean-square displacement (*MSD*) and the self-overlap function, *Q*(*t*), (defined in the Materials and Methods) as a function of time *t* show typical glassy behavior (Figs. 2b,c). We define relaxation time, *τ*, as *Q*(*t* = *τ*) = 0.3. For a particular *P*_0_, the glass transition temperature, *T*_*g*_, is defined as *τ* (*T*_*g*_) = 10^4^.

**FIG 2.**
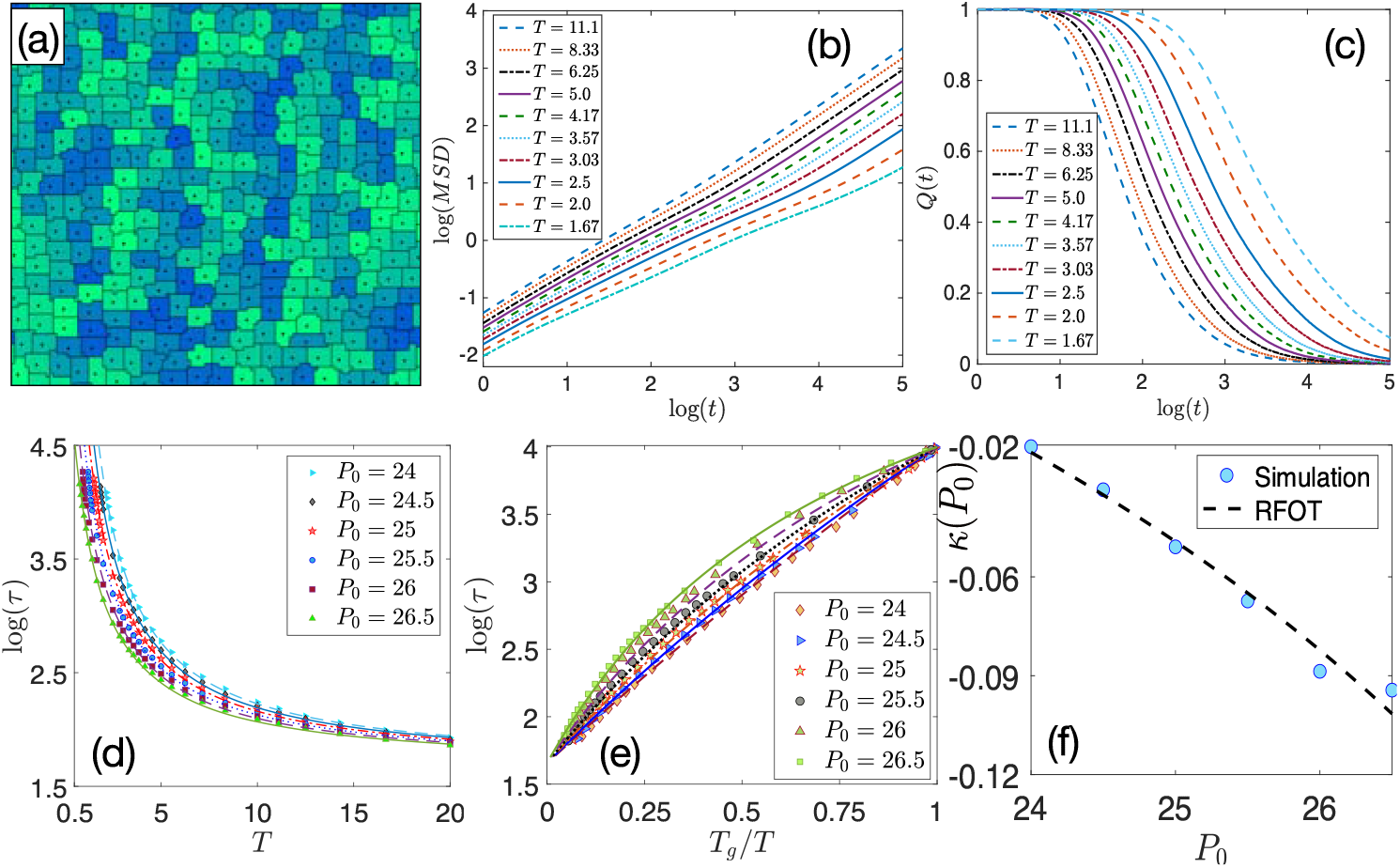
Behavior of CPM in the low-*P*_0_ regime. (a) Typical configuration of a system at *P*_0_ = 25 and *T* = 2.5, close to *T*_*g*_. Due to the underlying lattice structure, minimum perimeter configuration for a certain area is a square that shows up in the low *T* configuration. (b) Mean square displacement (MSD) and (c) self-overlap function, *Q*(*t*), as a function of time *t* for *P*_0_ = 25 show typical glassy behaviors where growth of MSD and decay of *Q*(*t*) become slower with decreasing *T*. (d) Relaxation time *τ* as a function of *T* for different *P*_0_, symbols are simulation data and lines are the corresponding RFOT theory plots (Eq. 5). (e) Angell plot in this regime shows sub-Arrhenius relaxation, symbols are data and lines are RFOT theory predictions. (f) Simulation data (symbols) for kinetic fragility, *κ*(*P*_0_), in this regime also agree well with the RFOT theory prediction (line).

We now develop the RFOT theory for CPM in this regime. Our approach is perturbative in nature where we treat a confluent system with *P*_0_ = 23 as our reference system around which we expand the effect of *P*_0_ (see SM Sec. S3 for details). The perimeter constraint in the form of *P*_0_ is written as some interaction potential Φ(*P*_0_) and Φ(23) gives the reference system potential. Then *s*_*c*_[Φ(*P*_0_)] can be written as

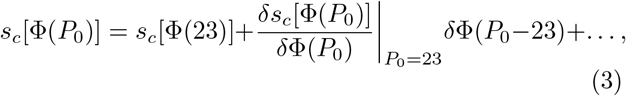

where 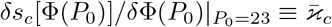 and *δ*Φ(*P*_0_ − 23) = *c*_1_(*P*_0_ − 23), with 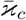 and *c*_1_ being constants. For the reference system, *s*_*c*_[Φ(23)] (*T* − *T*_*K*_), where *T*_*K*_ is the Kauzmann temperature [41, 42, 52, 54], and therefore, for CPM, we obtain *s*_*c*_ ∼ [*T* − *T*_*K*_ + ϰ_*c*_(*P*_0_ 23)] where 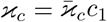. Similarly, for the surface reconfiguration energy, we can write

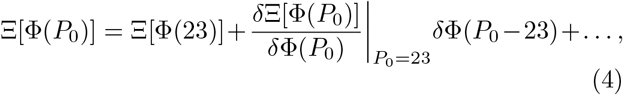

where 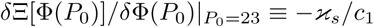 is constant. As *P*_0_ increases, the interaction potential decreases allowing more configurations available to the system and making it easier for reconfiguration, this increases *s*_*c*_ and reduces Ξ respectively, explaining the opposite signs for ϰ_*c*_ and ϰ_*s*_ in Eqs. (3) and (4). Writing *E*Ξ[Φ(23)] ≡ *k*_1_ and *E*ϰ_*s*_ *k*_2_, we obtain from Eq. (2) the RFOT theory expression for *τ* as

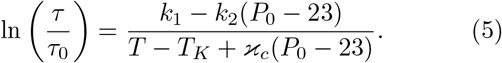

The constants *k*_1_, *k*_2_, *T*_*K*_ and ϰ_*c*_ are independent of *T* and *P*_0_; they only depend on the microscopic details of a system and dimension. For a given system, we treat these constants as fitting parameters in the theory and obtain their values from fit with simulation data. Note that *τ*_0_ depends on the high *T* properties of the system, which is nontrivial and will be explored elsewhere. Our analysis in the low-*P*_0_ regime shows that *P*_0_-dependence of *τ*_0_ is weaker and can be taken as a constant since our interest in this work is the glassy regime.

The minimum possible perimeter in our simulation is 26 (Materials and Methods) and we expect the critical *P*_0_ separating the two regimes to be somewhere between 27 and 28. We first concentrate on the results for *P*_0_ = 24 to 26.5 as this is, possibly, the most relevant regime experimentally and present *τ* as a function of *T* for different *P*_0_ in Fig. 2(d). We fit one set of data presented in Fig. 2(d) with Eq. (5) and obtain the parameters as follows: *τ*_0_ = 45.13, *k*_1_ = 14.78, *k*_2_ = 1.21, *T*_*K*_ = 0.0057 and ϰ_*c*_ = 0.31. Note that with these constants fixed, there is no other fitting parameter in the theory, we now show the plot of Eq. (5), as a function of *T* for different values of *P*_0_ with lines in Fig. 2(d). Figure 2(e) shows the same data in Angell plot representation that shows *τ* as a function of *T*_*g*_/*T* in semi-log scale. All the curves meet at *T* = *T*_*g*_ by definition. The simulation data agree well with RFOT predictions at low *T* where the theory is applicable.

One striking feature of the Angell plot in Fig. 2(e) is the sub-Arrhenius nature of *τ* whereas it is super-Arrhenius in most glassy systems. Similar results were reported for voronoi model in Ref. [26] demonstrating similarities between CPM and vertex-based models. Within our RFOT theory, the sub-Arrhenius relaxation appears due to the distinctive interaction potential imposed by the perimeter constraint, in a regime controlled by geometric restriction, and appears when the system is about to satisfy the perimeter constraint. An important characteristic of this regime is that [*T*_*K*_ − ϰ_*c*_(*P*_0_ − 23)] becomes negative. We get back to this point later in the paper when we subject our RFOT theory to more stringent tests.

One can define a kinetic fragility, *κ*(*P*_0_), and fit the simulation data for different *P*_0_ with the form 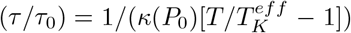. We present *κ*(*P*_0_) in Fig. 2(f) where symbols are values obtained from fits with simulation data and the dotted line is theoretical prediction, the agreement, again, is remarkable. Fragility of the system decreases as *P*_0_ increases and *κ*(*P*_0_) becomes more negative consistent with stronger sub-Arrhenius behavior.

### Large-*P*_0_ regime

In this regime most cells satisfy the perimeter constraint, cell boundaries are nonlinear [Fig. 3(a)], and dynamics becomes independent of *P*_0_ implying constant values of Ξ and *s*_*c*_. Then the RFOT theory becomes

**FIG 3.**
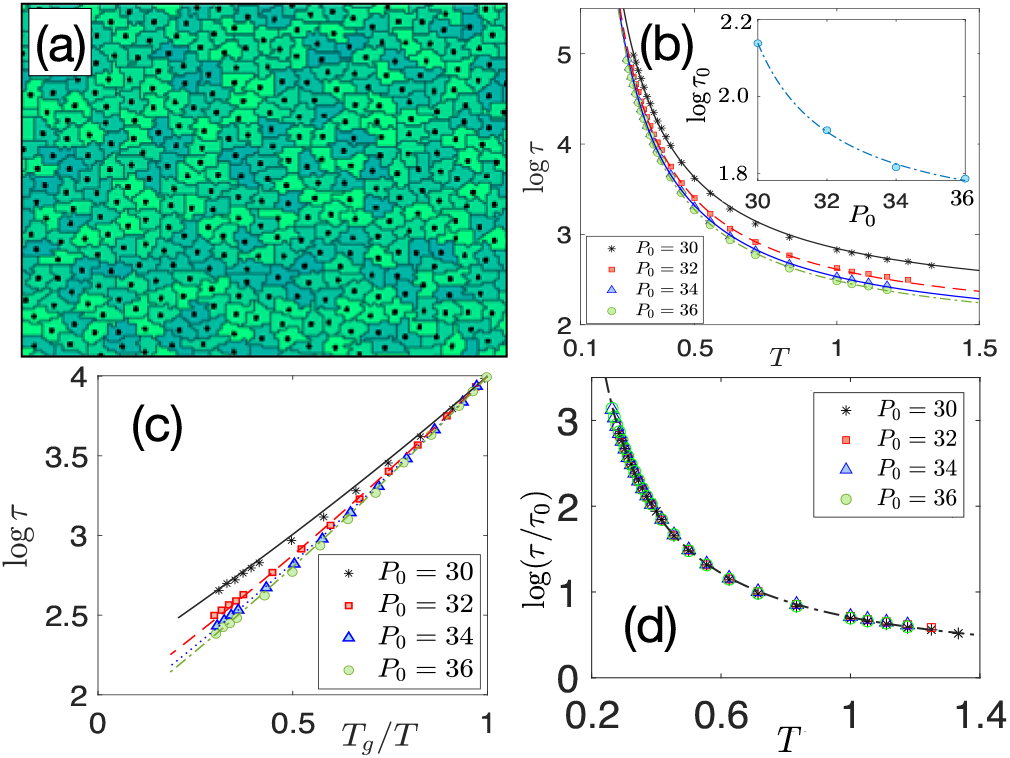
Behavior of CPM in the large-*P*_0_ regime. (a) Typical configuration of the system with *P*_0_ = 34 and *T* = 0.5 close to *T*_*g*_. The cell boundary assumes a fractal-like nature to satisfy the perimeter constraint. (b) Relaxation time *τ* as a function of *T*, symbols are data and lines are RFOT theory fits (Eq. 6). **Inset:** *τ*_0_ as a function of *P*_0_. The line is a fit with a function *τ*_0_(*P*_0_) = *a* + *b*/(*P*_0_ − *c*) with *a* = 3.79, *b* = 2.58 and *c* = 27.72. (c) Angell plot for the same data (symbols) as in (b), lines are RFOT theory plots (Eq. 6). (d) *τ*/*τ*_0_ for different *P*_0_ follow a master curve, the line is RFOT theory result. The data collapse illustrates that glassiness in this regime is independent of *P*_0_.

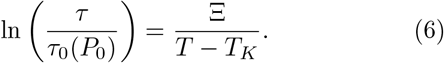

Although *P*_*i*_ = *P*_0_ on the average, there are fluctuations of *P*_*i*_ around *P*_0_ when *T* ≠ 0. These fluctuations govern the interaction potential and are stronger at higher *T*. Thus, *P*_0_-dependence of *τ*_0_ is important in this regime.

Figure 3(b) shows *τ* as a function of *T*; they clearly vary for different *P*_0_ and this difference comes from *P*_0_- dependence of *τ*_0_. We fit Eq. (6) with one set of data and obtain Ξ = 1.54, *T*_*K*_ = 0.052 and a corresponding value for *τ*_0_(*P*_0_). Keeping Ξ and *T*_*K*_ fixed, we next fit rest of the data to obtain *τ*_0_(*P*_0_). The fits are shown by lines in Fig. 3(b) and *τ*_0_(*P*_0_) is shown in the inset where the line is a proposed form: *τ*_0_(*P*_0_) ∼ 1/(*P*_0_ − constant). Figure 3(c) shows the Angell plot representation of the same data as in (b) and the corresponding RFOT theory plots are shown by lines. Figure 3(d) shows *τ*/*τ*_0_ as a function of *T* for different values of *P*_0_, all the data following a master curve support our hypothesis that *P*_0_-dependence in this regime comes from *τ*_0_(*P*_0_). More important, one would expect no glassy behavior in this regime if rigidity transition controlled glassiness as in the vertex models [23, 24]. In contrast, CPM shows the presence of glassy behaviour even in this regime and the existence of disordered configurations implies it is also expected within RFOT theory. *q* at *T*_*g*_ in this regime is proportional to *P*_0_ [Fig. 1(c)], thus, *q* cannot be an order parameter for glass transition.

### Further tests for extended RFOT theory

Having demonstrated that our RFOT theory captures the key characteristics of glassiness in a confluent system, we now subject our theory to stringent tests through three different questions:

Within the theory sub-Arrhenius behavior is found when the effective *T*_*K*_ is negative, i.e., *T*_*K*_ − ϰ_*c*_(*P*_0_ − 23) < 0 (Eq. (5)). This implies super-Arrhenius behavior for *P*_0_ ≤ (*T*_*K*_ + 23ϰ_*c*_)/ ϰ_*c*_ ≈ 23. We now simulate the system in this regime and show the Angell plot in Fig. 4(a) where symbols represent simulation data and the corresponding lines are RFOT theory predictions. We emphasize that these curves are *not* fits, we simply plot Eq. (5) with the constants as obtained earlier. All the relaxation curves for different *P*_0_ are super-Arrhenius as predicted by the theory. Figure 4(b) shows comparison of *T*_*g*_, obtained from simulation and RFOT theory, for different *P*_0_.

**FIG 4.**
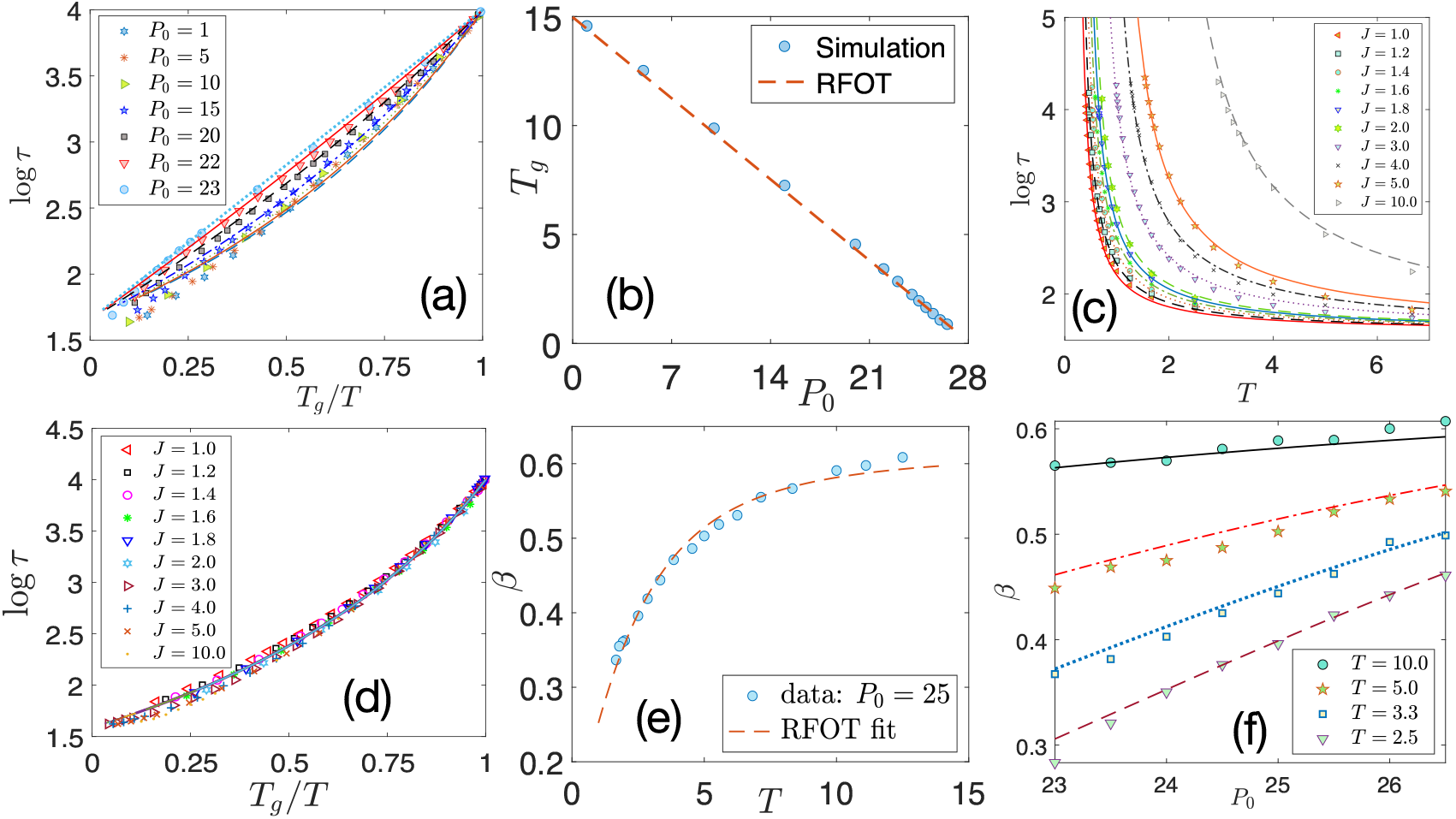
Tests of our extended RFOT theory. (a) Theory predicts super-Arrhenius behavior for *P*_0_ ≤ 23. Angell plot for low-*P*_0_ simulation data (symbols) agree well with the RFOT theory predictions, Eq. (5) (lines). (b) Comparison of *T*_*g*_ at different *P*_0_ between simulation (symbols) and RFOT theory (dashed line). (c) *τ* for the model with *λ*_*P*_ = 0 and different values of *J*, symbols are simulation data and lines are RFOT theory (SM Eq. S18). (d) Our theory predicts super-Arrhenius behavior and constant fragility when *λ*_*P*_ = 0, this is well-supported by simulation data (symbols) that follow a master curve in the Angell plot representation, the lines are RFOT theory plots. (e) Stretching exponent *β* for *P*_0_ = 25 as a function of *T*. Fit of simulation data with the RFOT theory expression, Eq. (7), gives 𝒜 = 0.62 and ℬ = 0.3. (f) Trends of *β* as a function of *P*_0_ at different *T* agree well with the RFOT theory predictions, Eq. (7), with 𝒜 and ℬ obtained from the fit in (e).

Next, our theory traces the sub-Arrhenius behavior and negative kinetic fragility to the perimeter constraint in Eq. (1). Hence, if we set *λ*_*P*_ = 0 and look at the glassy behavior as a function of *J* (Eq. 1), the system should show not only super-Arrhenius behavior, but also a constant fragility (i.e., a master curve in Angell plot). Figure 4(c) shows simulation data for *τ* as a function of *T* for different *J* (symbols) as well as the corresponding RFOT theory (SM, Eq. S18) predictions (lines). The Angell plot corresponding to these data are shown in Fig. 4(d). Indeed, we find that this system exhibits super-Arrhenius relaxation and data for different *J* follow a master curve, in agreement with our theory. These results are important from at least two aspects: first, they show the predictive power and applicability of the theory, second, systems with and without the perimeter constraint are qualitatively different [39].

Finally, we compare the stretching exponent *β* [55, 56] that describes decay of the overlap function *Q*(*t*) ∼ exp[(−*t*/*τ*)^*β*^]. The RFOT expression for *β* (SM, Sec. S4) is

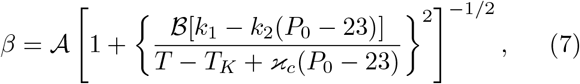

where 𝒜 and ℬ are two constants; we fit Eq. (7) with the simulation data for *P*_0_ = 25, as shown in Fig. 4(e), and obtain 𝒜 = 0.62 and ℬ = 0.3. We then compare the RFOT predictions with simulation data for different *P*_0_ as shown in Fig. 4(f) for four different *T*. Again, the trends for *β* agree quite well with theoretical predictions.

## II. DISCUSSION AND CONCLUSION

Complete confluency imposes a strong geometric restriction bringing about two different regimes as *P*_0_ is varied. Our theory traces the unusual sub-Arrhenius behavior to the distinctive nature of interaction potential resulting via the perimeter constraint and shows up in a regime where the system is about to satisfy this constraint. Qualitative similarities of the results presented here with those from vertex-based simulations [23, 26] seem to suggest glassiness in such systems depends on two key elements, first, the energy function, and second, the confluent nature, and *not* the microscopic details, of the models. We believe, the RFOT theory that we have developed is applicable to a general confluent system and not restricted to CPM. The three predictions of the theory that we have discussed, namely super-Arrhenius behavior in a different region of low-*P*_0_ regime, super-Arrhenius and constant fragility in a model with *λ*_*P*_ = 0 and the stretching exponents at different *P*_0_ agree well with simulation data. These predictions can easily be tested in vertex-based simulations, such results will further establish the similarity (or the lack of it) of such models with CPM.

Vertex-model simulations have argued the rigidity transition controls the glassy dynamics, and the observed shape index, *q*, has been interpreted as a structural order parameter for glass transition [14, 23, 24]. Our study shows these results are *not* generic for models of confluent systems. Within CPM, lowest value of *q* is determined by geometric restriction in the low-*P*_0_ regime whereas it is proportional to *P*_0_ in the large-*P*_0_ regime although glassiness is found in both and controlled by the underlying disordered configuration.

Control parameters of glassiness in a confluent system are different from those in particulate systems. The experiments of Ref. [11] on human mammary epithelial MCF-10A cells show that expression of RAB5A, that does not affect number density, fluidizes the system. Careful measurements reveal RAB5A affects the junction proteins in cortex that determines the target perimeter *P*_0_ [11, 57], which is a control parameter for glassiness in such systems. Apart from biological importance, we believe, CPM provides an interesting system to study from purely theoretical point of view to understand glassy dynamics in a new light.

The quantitative values of the parameters are lattice-dependent due to the presence of microstructure within CPM, e.g., a square is the minimum perimeter to area configuration on square-lattice. However, the qualitative results, such as, increase in inter-cellular repulsive interaction as *P*_0_ decreases, constant value of *q* in the low-*P*_0_ regime when the system is glassy, agree well with experiments as well as vertex model simulations [14]. A crucial difference between CPM and vertex model results lies in the mechanism that controls glassiness: the disordered energy landscape in the former, and a rigidity transition in the latter. A critical test will be to see if glassiness exists in the large-*P*_0_ regime in experiments along the lines of Refs. [57, 58]. The presence of metabolic activity in a biological system is effectively *T* in CPM. As metabolic activity reduces, self-propulsion, which is absent in the current model, also decreases. Therefore, presence or absence of glassiness in the large-*P*_0_ regime as metabolism is decreased will distinguish the two mechanisms.

## Materials and methods

### Simulation details

Details of the CPM and the Monte Carlo dynamics at temperature *T* are given in SM. For the results presented here, unless otherwise specified, we use a system of size 120 × 120 with 360 cells and an average cell area of 40. The minimum possible perimeter for a cell with area 40 on a square lattice is 26. We start with a rectangular cell initialization with 5 × 8 sites having same Potts variable and equilibrate the system for at least 8 × 10^5^ MC time steps before collecting data. We have set *λ*_*A*_ = 1 and *λ*_*P*_ = 0.5 for the results presented here. Data for other values of *λ*_*P*_, system sizes, and cell sizes are presented in the SM (Sec. S6).

### Mean square displacement and self-overlap function

Dynamics is quantified through the mean square displacement (*MSD*) and the self-overlap function, *Q*(*t*). *MSD* is defined as

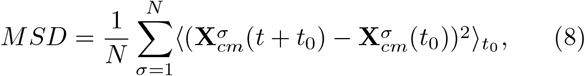

where 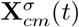 is center of mass of cell *σ* at time *t*, ⟨… ⟩_*t*_0__ denotes averaging over initial times *t*_0_ and the overline implies an averaging over ensembles. Unless otherwise stated, we have taken 50 *t*_0_ averaging and 20 configurations for ensemble averaging. *Q*(*t*) is defined as

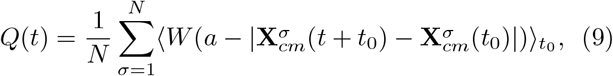

where *W* (*x*) is a heaviside step function

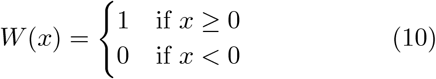

and *a* is a parameter that we set to 1.12.

## Acknowledgements

We thank Mustansir Barma and Chandan Dasgupta for many important and enlightening discussions and critical comments on the manuscript. We also thank Tamal Das, Kabir Ramola, Navdeep Rana, Kallol Paul and Pankaj Popli for discussions. This project was funded by intramural funds at TIFR Hyderabad from the Department of Atomic Energy (DAE), Government of India.

## Supplementary Material

In this Supplementary Material, we provide a brief description of the cellular Potts model, a discussion of the energy function, simulation details, a discussion of the contrasting roles of the target area and the target perimeter and an illustration of a *T*1 transition that is naturally included within the model in Sec. S1 followed by a detailed discussion on the source of metastability in the model and the nature of dynamics in Sec. S2. Section S3 provides detailed calculation of our extended random first order transition theory and the calculation of the stretching exponent is presented in Sec. S4. Sections S5-S8 provide results for the dynamics in the large-*P*_0_ regime for odd values of *P*_0_, results for different *λ*_*P*_, effects of finite system sizes and effects of different cell sizes on the dynamics respectively, followed by a brief discussion on the quantitative values of perimeter in CPM due to the underlying lattice structure in Sec. S9.

### S1. CELLULAR POTTS MODEL FOR BIOLOGICAL TISSUES

The cellular Potts model (CPM) is a mathematical, stochastic, computational lattice based model to simulate the behavior of cellular systems [1–3]. This model is also known as “extended large-*q* Potts model” and the “Glazier-Graner-Hogeweg (GGH) model” [3–5]. CPM can be simulated both for a single cell as well as a collection of cells with or without fluid in any spatial dimension *d*. It has applications in embryogenesis [5], cell sorting [3, 4], gradient sensing [2], wound healing [6], and many others. Since our interest in this work is the dynamics of a densely packed cellular monolayer, we restrict our discussion in 2*D*.

**FIG S1.**
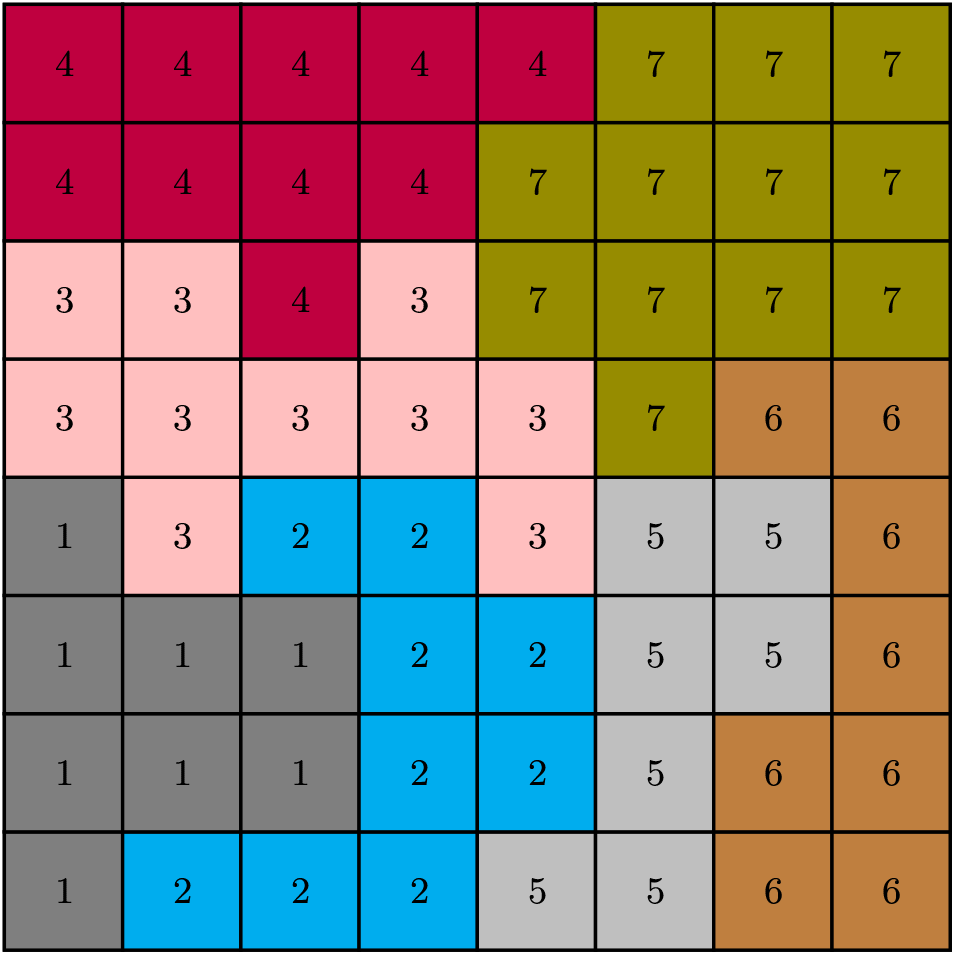
Cell representation in two dimensional CPM. Each of the lattice sites with the same cell index (denoted by a number and a color) belongs to a particular cell.

For the CPM in 2*D*, we use a square lattice of size *L* × *L* to represent a confluent cell monolayer. Each cell in this lattice consists of a set of lattice sites with same integer Potts spin (*σ*), also known as cell index, where *σ* ∈ [0, *N*], *N* being total number of cells. *σ* = 0 is usually reserved for fluid that is absent in our model. A typical lattice structure of cells in a 2*D* CPM is presented in Fig. (S1).

The cells in this model are evolved by stochastically updating one lattice site at a time through Monte Carlo (MC) simulation via an effective energy function ℋ, given by

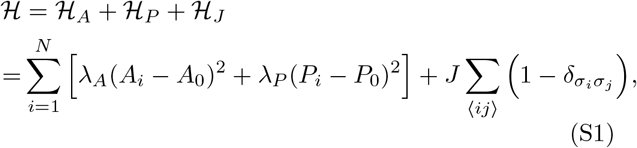

where ⟨*ij*⟩ signifies nearest neighbors, *A*_*i*_ and *P*_*i*_ are the area and the perimeter of *i*^*th*^ cell in the monolayer. *A*_0_ and *P*_0_ are the target area and the target perimeter of the cells, chosen to be the same for all cells in a monodisperse system. *λ*_*A*_ and *λ*_*P*_ are the elastic constants associated with the area and the perimeter constraints respectively. Interaction between two cells *σ*_*i*_ and *σ*_*j*_ is given by the parameter *J*, chosen to be same for all cells. *δ*_*ij*_ is the usual “Kronecker delta function” with the following property: *δ*_*ij*_ = 1 if *i* = *j* and 0 otherwise. *δ*_*ij*_ ensures that the contribution due to the third term in Eq. (S1) is zero when the sites are of same types, thus *ℋ*_*J*_ is proportional to *P*_*i*_ for *i*th cell. Positive values of *J* signify inter-cellular repulsion, whereas negative values give attractive interaction.

#### S1A. Significance of different terms in Eq. (S1)

Cells can be treated as incompressible in 3*D* [43]. It has been found in experiments that the height of a monolayer remains almost constant [21]. These two findings together allows a 2*D* description of the system with an area constraint written as ℋ _*A*_ in Eq. (S1). On the other hand, mechanical properties of a cell is mostly governed by cellular cortex [43] and this can be encoded in a perimeter constraint in the form of ℋ _*P*_. Inter-cellular interactions through different junction proteins like E-Cadherins and effects of pressure, contractility, cell adhesion etc are included within an effective interaction term, ℋ_*J*_ [21].

**FIG S2.**
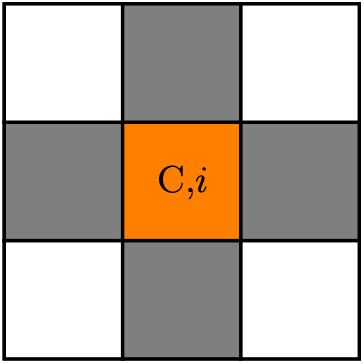
Von Neumann Neighborhood in CPM: Von Neumann neighbors of the central (C) site at *i* with color orange are the sites in color gray.

Note that the proteins affecting the inter-cellular interactions also affect the cortical properties, this implies the possibility of including ℋ_*J*_ in the perimeter constraint term, ℋ_*P*_ in Eq. (S1). As discussed in the previous section, ℋ_*I*_ for the *i*th cell is proportional to its perimeter: 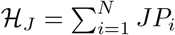. Therefore, we can rewrite Eq. (S1) as,

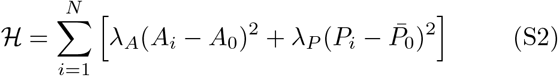

where 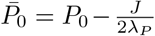 is the scaled target perimeter. Note that we have neglected a constant in Eq. (S2) since it does not affect the behavior of the system. For a particular *λ*_*P*_, dynamics of the system remains unchanged if we vary *P*_0_ and *J* in such a way that 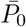 remains same.

In the simulations, we always set *λ*_*A*_ = 1 and then *λ*_*P*_ and *P*_0_ are the two control parameters. We have explored the dynamics for three different values of *λ*_*P*_ and as a function of *P*_0_.

#### S1B. Simulation Details

Dynamics at a temperature *T* within CPM proceeds through the stochastic attempts to update the cell index of a randomly chosen lattice site at each step; unit of time is defined by *L*^2^ attempts of such elemental steps, which are described below:

1. We randomly chose a candidate site *i* and a target site *j* from the nearest neighbors of *i*. Cell indices at *i* and *j* are *σ*_*i*_ and *σ*_*j*_.
2. Dynamics proceeds further only if *σ*_*i*_ ≠ *σ*_*j*_.
3. Check the local connectivity (see below) of the cells if the update is accepted, dynamics proceeds only if local connectivity remains intact.
4. Evaluate the change in energy, Δℋ, if *σ*_*i*_ is updated with *σ*_*j*_.
5. Accept the move with a probability *P*(*σ*_*i*_ → *σ*_*j*_) = min(1, *e*^−Δ*ℋ*/*T*^), where we have set Boltzmann constant *k*_*B*_ to unity.

During evolution, a site can only be updated with one of it’s Von Neumann Neighbors (VNN) defined in Fig. S2. If a cell is not locally simply connected, we designate it as a “fragmented cell”. In our simulation, we have followed the “Connectivity Algorithm (CA)” developed by Durand and Guesnet [7] and ensure local connectivity of the cells at all times. Centers of mass of the cells are calculated via the algorithm developed by Bai and Breen [8]. Cell division and apoptosis [9, 10] are forbidden in our simulation as we are interested in the equilibrium properties.

For most of the results presented in the main text, we have chosen a square lattice of size 120 × 120 with 360 total number of cells in the system. Average area of cells in the system is 40 and the minimum possible perimeter on a square lattice with this area is 26. We have kept *λ*_*A*_ = 1 fixed and simulated the system for different values of *λ*_*P*_. *λ*_*P*_ = 0.5 for the results presented in the main text and other values of *λ*_*P*_ are presented below in Sec. S6. Note that the dynamics is independent of *A*_0_ in our system (Sec. S1 S1C), we have simply used *A*_0_ as the average cell area such that the energy contribution from the area term for a cell with average area is zero; this is akin to adding a constant to the energy function.

We first equilibrate our system for 8 × 10^5^ MC time steps before the start of collecting data for mean-square displacement (*MSD*) and self-overlap function, *Q*(*t*) (defined in the materials and methods). Unless otherwise stated, each of the results is an average over 50 initial times *t*_0_ and 20 ensembles. To test system size effect, we have also studied systems with different simulation box sizes with the largest being 200 × 200 having 1000 cells with average cell size 40 and to observe the finite size effect, we have studied systems with simulation box size 180 × 180 and cell size 10 × 9. For average inherent structure energy calculation (Sec. S2), we first equilibrate the system at *T* = 4.0 for 7 × 10^4^ time steps and then decrease *T* to 0 in steps of 0.5 every 2000 time steps; we have also studied other equilibration *T*, equilibration times and rates to ensure expected behavior. To observe the system size effect in 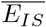, we have taken simulation box of sizes, 200 × 200, 160 × 160, 120 × 120 and 80 × 80 for *P*_0_ = 25; for this data, we have used at least 1000 ensembles and larger number of ensembles for smaller systems to ensure good averaging.

The most time consuming part in the simulation is calculation of cellular perimeter at each elemental step. We have developed an algorithm [11] that allows us to locally calculate the perimeter and makes it possible to simulate large system sizes for long times, appropriate to investigate the glassy dynamics.

**FIG S3.**
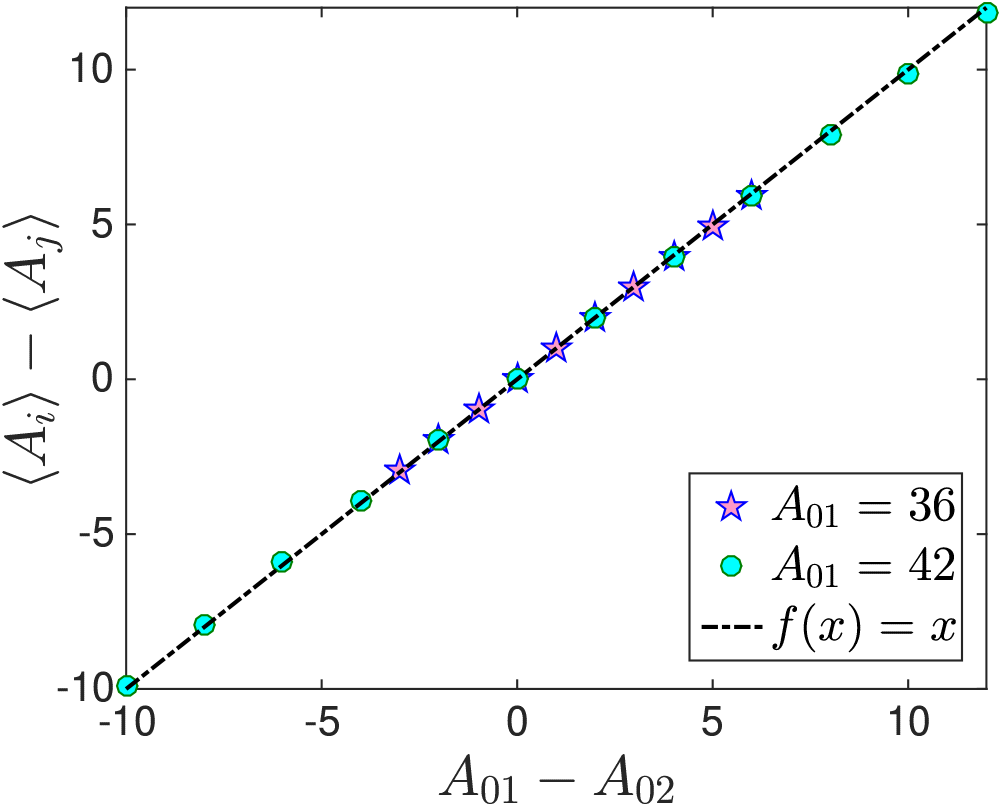
We have chosen a binary system with different values of target areas. ⟨*A*_*i*_ ⟩ and ⟨*A*_*j*_ ⟩ correspond to the average areas of cells with target areas *A*_01_ and *A*_02_ respectively. Eq. (S6) predicts ⟨*A*_*i*_ ⟩ − ⟨*A*_*j*_ ⟩ should be equal to (*A*_01_ − *A*_02_) and this is supported by the simulation data. We have chosen a system of 160 cells with system size 80 × 80 and each point is averaged over 10^3^ *t*_0_ values.

#### S1C. Discussion on *A*_0_ and *P*_0_ in the energy function

We have shown in the main text that the target area in Eq. (S1) does not affect the dynamics, whereas the target perimeter *P*_0_ plays the role of a control parameter. Let us first look at the change in energy, Δℋ _*area*_, for *σ*_*i*_ → *σ*_*j*_ in a monodisperse system coming from the area term alone,

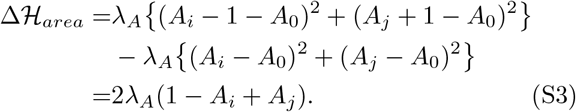

Thus, Δ ℋ _*area*_ is independent of *A*_0_ for a monodisperse system and thus, *A*_0_ can not affect the dynamics. Let us now consider dispersion of *A*_0_, and for simplicity, we consider a binary system with two different target areas *A*_01_ and *A*_02_. At each of the elemental steps, when candidate cell and target cell have same target areas, the situation becomes similar as depicted in Eq. (S3), where *A*_0_ does not affect the dynamics. Let us then examine the case where the two cells have different target areas, *A*_01_ for the *i*th cell and *A*_02_ for the *j*th cell.

There can be two scenarios in this case, either the target areas are consistent with the condition of complete confluency or they are not. In the first case, average areas of cells will be given by their respective target areas and the individual cell areas can be written as *A*_*i*_ = *A*_01_ +*δA*_*i*_ and *A*_*j*_ = *A*_02_ + *δA*_*j*_, where *δA*_*i*_ and *δA*_*j*_ are fluctuations from their average values (see the derivation below). The change in energy coming from the area term for the attempted MC move then becomes

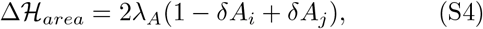

and thus, independent of *A*_01_ and *A*_02_. On the other hand, when the target areas are not consistent with the constraint of complete confluency, average area is still set by them. Since each elemental MC step consists of two cells, it suffices to consider a system of two cells, the area part of the energy function is

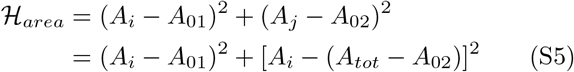

where *A*_*tot*_ = *A*_*i*_ + *A*_*j*_ is the total area. Minimizing Eq. (S5), we obtain the average cell areas as

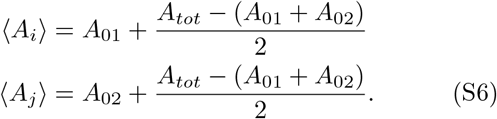

Note that the second term in the right hand side in Eq. (S6) becomes zero when target areas are consistent with constraint of confluency, that is, (*A*_01_ + *A*_02_) = *A*_*tot*_. Thus, we expect the difference of average areas of the two cells to be *A*_01_ − *A*_02_ and this agrees with simulation data, as shown in Fig. S3. Going through a similar argument as above, we obtain Δ ℋ _*area*_ to be same as in Eq. (S4) in this case as well. This argument can also be extended for a polydisperse system. Thus, we see that in a confluent monolayer, target area of the cells cannot affect the dynamics. Note that {*A*_0*i*_}, however, has a meaning for the statics of the system. It determines the average cell areas of the different types of cells. Average cell area is a geometric quantity that determines the critical value of *P*_0_ separating the two regimes in the dynamics.

We now look at the role of *P*_0_ and for this we discuss the perimeter term alone. Let us consider two values of target perimeter, 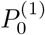 and 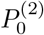 such that 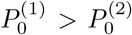. For the perimeter term, we can write

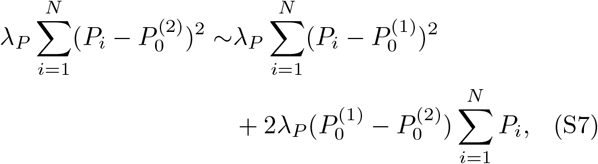

where we have ignored the constant part. Since 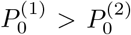, the linear term in the right hand side represents repulsive interaction. This result illustrates that decreasing *P*_0_ results in increased repulsive interaction, consistent with the experiments of Park *et al* [12]. Thus, *P*_0_ can be thought of as a parameter of intercellular interaction and hence works as the control parameter for the dynamics. Eq. (S7) also shows that the change in potential due to change in target perimeter, *δP*_0_, is linearly proportional to *δP*_0_. The source of the contrasting roles of *A*_0_ and *P*_0_ is the condition of complete confluency in the system: change in *A*_*i*_ of the two cells in an elemental MC step must be same to ensure the system remains confluent, whereas that in *P*_*i*_ need not be the same.

**FIG S4.**
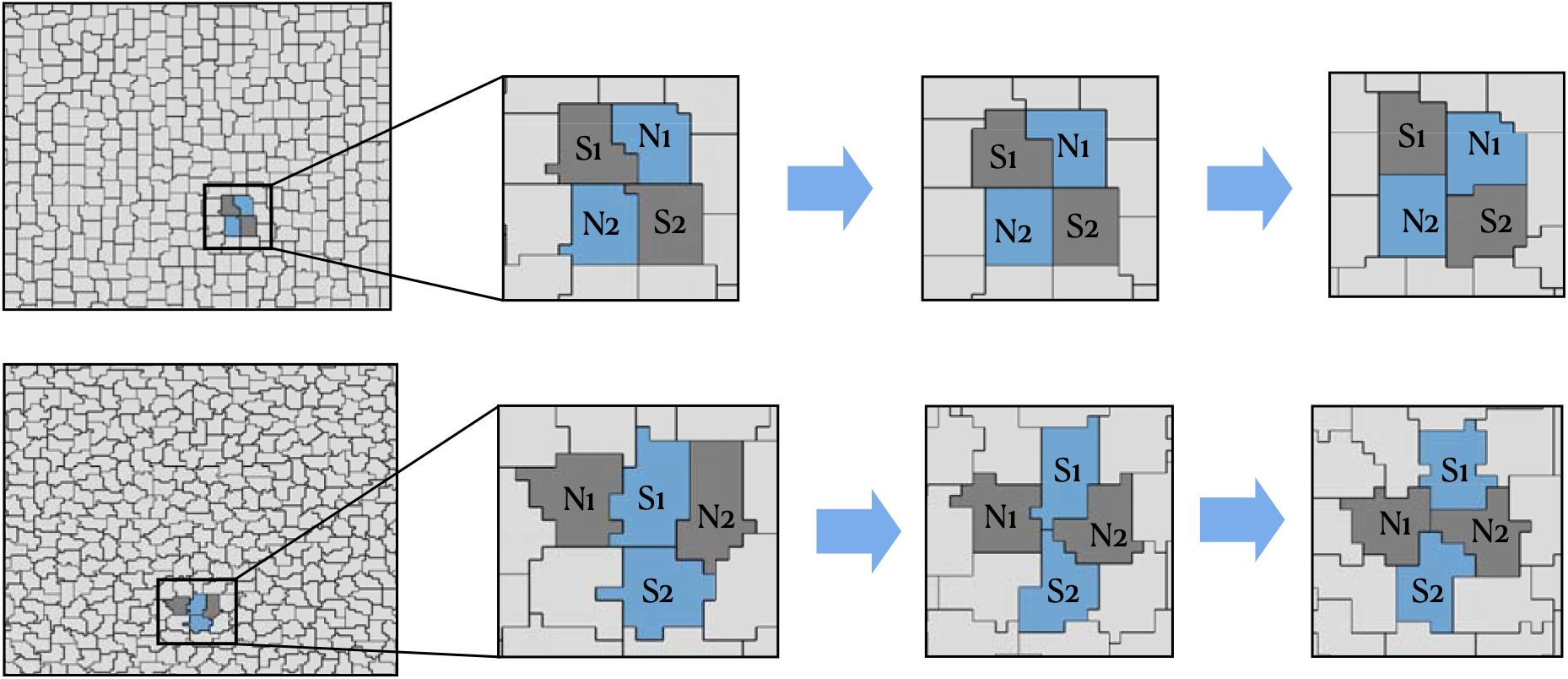
Snapshots of neighbor exchange or a *T*1 transition process in CPM. Upper panel shows a *T*1 transition event for *P*_0_ = 25 at *T* = 2.0 and the lower panel for *P*_0_ = 32 at *T* = 0.5; we follow the time evolution of four cells shown by the marked regions in the system (left most figures) and show the configurations of these cells at three different times. At the first snapshots, *S*_1_ and *S*_2_ share a common boundary whereas *N*_1_ and *N*_2_ don’t. The scenario reverses in the third snapshot.

#### S1D. *T*1 Transition

Dynamics in a biological tissue proceeds via a series of complicated biochemical processes that are simply represented via an effective temperature *T* within CPM. At the coarse grained level, an important process for dynamics is known as the *T*1 transition where cells exchange their neighbors [13]. Within vertex-based models, *T*1 transitions need to be carefully implemented with a certain rate. Since *T*1 transitions are the only mode of dynamics, it is then natural that the rate of such transitions is going to crucially affect the dynamics [14]. It remains unclear what controls this rate and how to connect it with different parameters in an equilibrium model, in the absence of activity, making it difficult for quantitative theoretical development. As discussed in the main text, *T*1 transitions are naturally included within CPM.

In the CPM simulation, as we allow cells to evolve, neighbor exchange and the junctional rearrangements take place via *T*1 transition, where a cell boundary between two cells shrinks to zero and a new cell boundary forms between two other cells that were initially not sharing common boundary. We show two such *T*1 transition processes from our simulation in Fig. (S4) for *P*_0_ = 25 at *T* = 2.0 (upper panel) and *P*_0_ = 32 at *T* = 0.5 (lower panel). In both panels, cells *S*_1_ and *S*_2_ share common boundary in the first snapshot, whereas *N*_1_ and *N*_2_ do not have any common boundary. As time progresses, the boundary between *S*_1_ and *S*_2_ shrinks (the middle figures) and eventually a common boundary between *N*_1_ and *N*_2_ forms whereas *S*_1_ and *S*_2_ depart from each other (last snapshots in Fig. S4). The rate of *T*1 transitions within CPM depends on *T* and *P*_0_, however, the quantitative details of how this rate compares with the rate in vertex-based models remains an important open question.

### S2. SOURCES OF METASTABILITY LEADING TO GLASSY DYNAMICS

It is hard to avoid crystallization in a monodisperse system of point particles in two dimension, but, as we show below, a confluent cellular system is different. The existence of metastability in Potts model in the limit of large Potts variable, greater than 4, is well-known [15], however, the nature and origin of metastability within CPM remains unclear. Hexagons can entirely tile space in 2*D* and the constraint of complete confluency makes polygons with six neighbors favorable. Nevertheless, there are many possible ways to completely tile space with polygons, and a distribution of different polygons is found in experiments [16]. Glass transition is driven by metastable configurations with a disordered energy landscape; it is crucial to understand the source of this metastability for a complete characterization of the glassy dynamics. We have shown in the main text that even in a monodisperse system, there exists a distribution of *A*_*i*_ and *P*_*i*_ in the steady state [Fig. 1(a)] effectively leading to a polydisperse system and allowing the system to easily avoid the periodic minimum. Figure S5(a) shows the instantaneous area and perimeter of a typical cell in the steady state: they fluctuate over time and the two do not necessarily follow each other, which is not surprising since it is possible to change one without changing the other.

**FIG S5.**
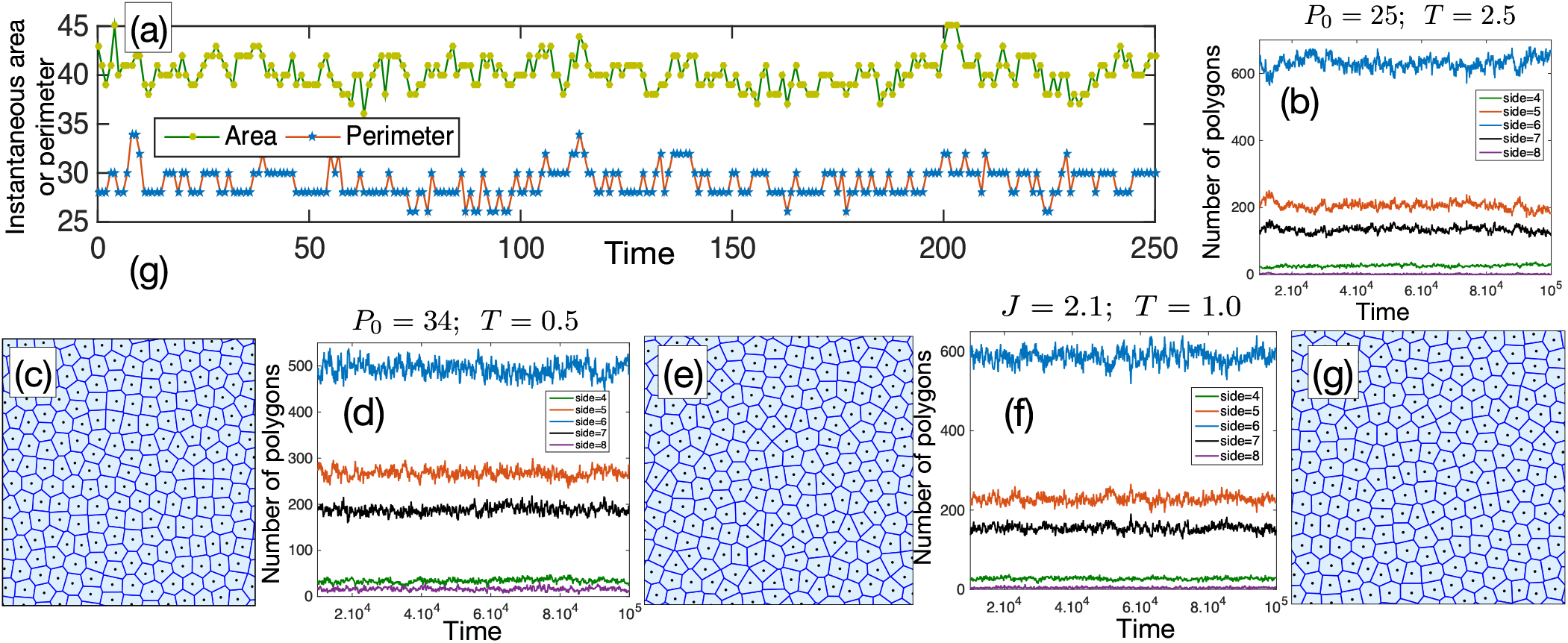
Instantaneous area and perimeter and distribution of polygons obtained through voronoi tessellation. (a) Variation of instantaneous area and perimeter of a typical cell in the system in steady state with *P*_0_ = 25 and *T* = 5.0. Zero of time, noted in the *x*-axis, is the point when we start collecting data. Note that fluctuation in area is unrelated to that of perimeter (see text). (b-g) Average number of polygons (irrespective of their regularity) in the steady state remains constant over time, signifying the topological disorder does not get annealed out over time. (b) and (d) for systems with *λ*_*P*_ = 0.5 and values of *P*_0_ and *T* as noted in the figures, (f) for a system with *λ*_*P*_ = 0, *J* = 2.1 and *T* = 1.0. (c), (e) and (g) are voronoi tessellations for the centers of mass of the cells in a typical configuration for the systems in (b), (d) and (f) respectively.

We now look at the topological nature of the disorder. A histogram of polygons (irrespective of their regularity), obtained via voronoi tessellation of centers of mass of the cells, reveals largest number of hexagons in steady state, as expected for a confluent system and shown in Figs. S5 (b) and (d) for two different values of *P*_0_ = 25 and 34 respectively at a *T* close to their *T*_*g*_. Figure S5(f) shows similar data for a system with *λ*_*P*_ = 0 and *J* = 2.1 at *T* = 1.0. Figures S5(c), (e) and (g) show the voronoi tessellation for centers of mass of a typical configuration of the systems in (b), (d) and (f) respectively. Figure S5 also shows the presence of a significant number of other polygons (mostly with sides 5 and 7) in steady state, implying presence of defects leading to disorder and that the system is trapped in a metastable minima.

Key to the random first order transition (RFOT) theory of glassy dynamics is a disordered energy landscape with extensive number of minima. As the system explores the energy landscape, it remains stuck in the metastable minima longer as *T* decreases leading to the slow dynamics. It is imperative to study the nature of the energy landscape of a system to infer applicability of RFOT theory. One straightforward quantity to study in this context is the configurational entropy, *s*_*c*_ = (log 𝒩)/*N*, where 𝒩 is the number of such metastable minima. Known methods of calculating *s*_*c*_ requires complete knowledge of a reference state which is taken as the high *T* liquid in particulate systems. However, the high *T* phase of a confluent system is highly nontrivial and not yet well-understood; thus, the conventional methods are not applicable. We investigate the role of metastability through another related quantity, the inherent structure energy, *E*_*IS*_, although the exact mathematical relation between *E*_*IS*_ and *s*_*c*_ is not yet known [17].

Within the inherent structure picture of glassy dynamics [18], only certain energy minima are accessible to the system at a particular *T*. If we equilibrate the system at a certain *T* and then set *T* = 0, the minimized energy, *E*_*IS*_, reflects the accessible energy minima at that *T*. We have shown in the main text that ensemble averaged inherent structure energy, 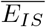, decreases linearly as *P*_0_ increases in the low-*P*_0_ regime and then it becomes zero for even values of *P*_0_ [Fig. 1(d)]. For odd values of *P*_0_ it goes to a different constant due to the residual interaction resulting from the fact that cellular perimeters in our system can only assume even values, however, the qualitative behavior remains same [Fig. S6(a)]: In the large-*P*_0_ regime, cells are able to satisfy both the area and the perimeter constraints when *P*_0_ is even. When *P*_0_ is odd, they satisfy the area constraint, but the perimeter fluctuates between *P*_0_ ± 1. We typically see a relatively larger number of cells have perimeter *P*_0_ + 1, this is expected as there is no restriction on larger perimeter with a certain area ^1^. Thus, the value of 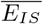 is either zero or *λ*_*P*_ *N* as shown in Fig. S6(a).

**FIG S6.**
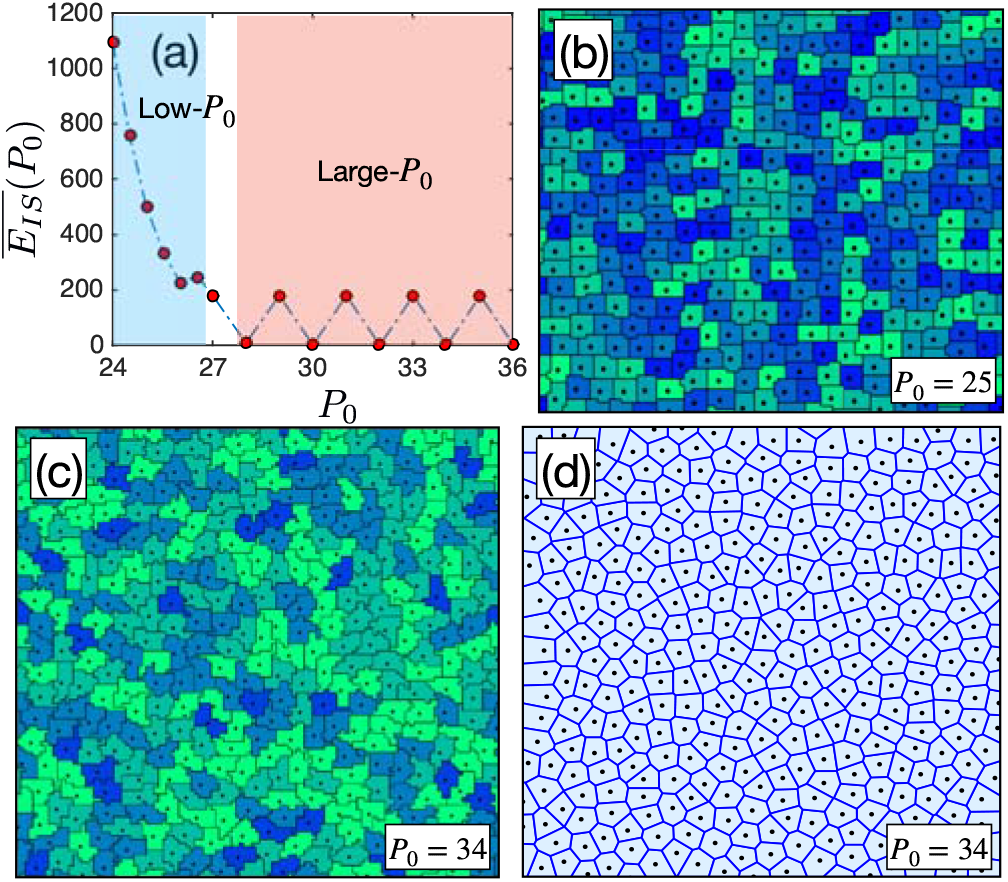
Inherent structure properties of CPM. (a) The ensemble averaged inherent structure energy, 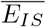, as a function of *P*_0_ has a strong *P*_0_-dependence in the low-*P*_0_ regime and then saturates to zero for even *P*_0_ and to *λ*_*P*_ *N* for odd *P*_0_ in the large-*P*_0_ regime (see text). (b) Inherent structure configuration for a system with *P*_0_ = 25. (c) Inherent structure configuration for a system with *P*_0_ = 34 in the large-*P*_0_ regime. Cells in this regime are able to satisfy the area and perimeter constraints when *P*_0_ is even, all the cells in the inherent structure of the system in this regime has average area and target perimeter, 40 and 34 in this case. (d) Voronoi tessellation of the centers of mass of the system shown in (c).

We show the inherent structure configuration for *P*_0_ = 25 in Fig. S6(b) and that for *P*_0_ = 34 in Fig. S6(c); the inherent structure configurations are indeed disordered and there exist many different configurations. The energy landscape in the large-*P*_0_ regime looks different from the one in the low-*P*_0_ regime; all the minima are at the same value of energy in the former whereas the energy for different minima are different in the latter. This leads to different behaviors in the two regimes. Note that, although energy of different minima are same in the large-*P*_0_ regime, they are separated by different energy barriers. As *T* decreases, it becomes difficult for the system to go from one minimum to another leading to the glassy dynamics as characterized in the main text. We also show the voronoi tessellation in Fig. S6(d) of the centers of mass of the inherent structure shown in Fig. S6(c) to emphasize that the actual perimeter is underestimated by a voronoi tessellation.

**FIG S7.**
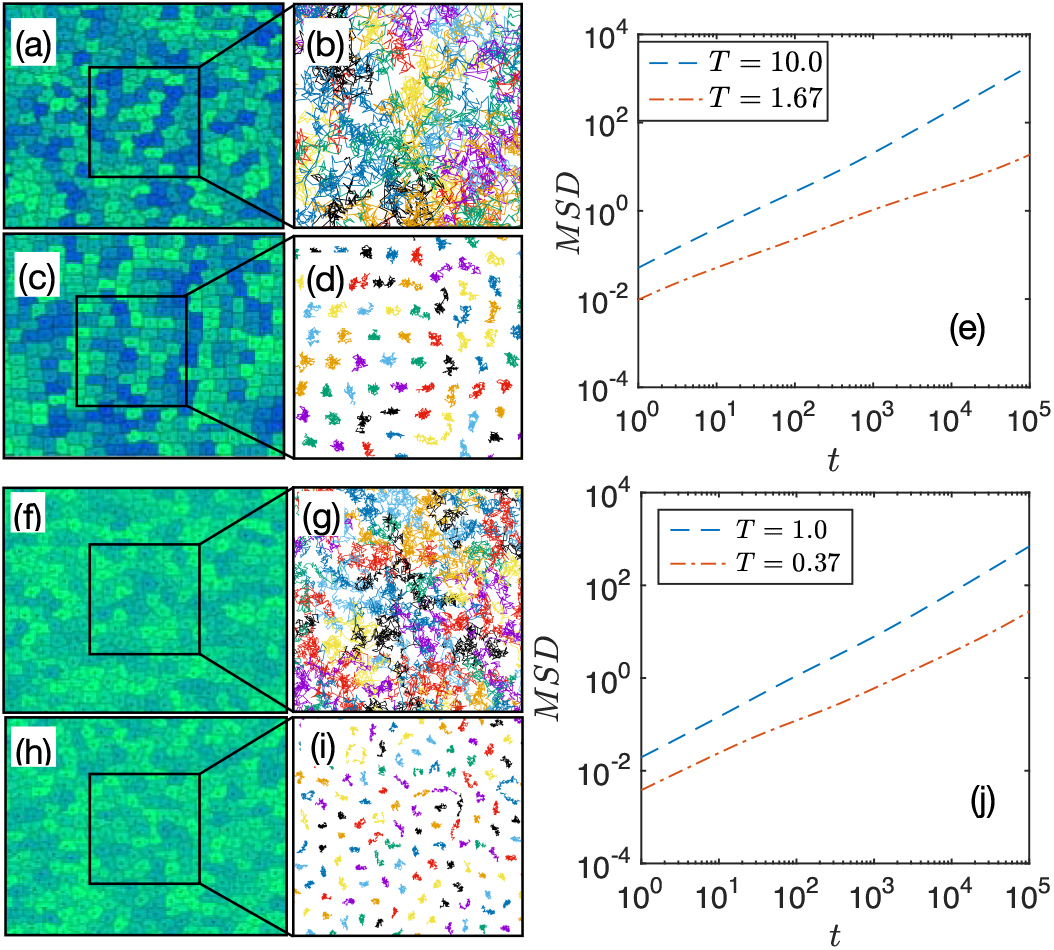
Qualitative nature of the dynamics. (a) A typical configuration for a system with *P*_0_ = 25 and *T* = 10. (b) Trajectories of the centers of mass of the cells inside a region as marked in (a). (c) and (d) are same as in (a) and (b) but at *T* = 1.67, close to the glass transition temperature *T*_*g*_ of the system. (e) Mean square displacement (*MSD*) of the system at the two different *T*. (f-j) Same plots as in (a-e) but for *P*_0_ = 32, the two different *T* chosen for these plots are *T* = 1.0 and *T* = 0.37 as shown in (j).

#### S2A. Qualitative nature of the dynamics

The system shows glassy behaviors in both low-*P*_0_ and large-*P*_0_ regimes although the behaviors in the two regimes are different due to geometric restrictions in the ability of the cells to satisfy the perimeter constraint. We show typical snapshots of the system for *P*_0_ = 25 and *P*_0_ = 32 in Fig. S7 at a high *T* (a and f) as well as at a low *T* close to *T*_*g*_ (c and h). We follow the trajectories of the centers of mass of the cells inside the regions marked in Fig. S7. At high *T*, the cells move quite a lot as revealed by the trajectories of their centers of mass in Figs. S7(b) and (g). As *T* decreases, the movements become small as seen in the trajectory plots in Figs. S7(d) and (i). The corresponding mean-square displacements (*MSD*) are shown in Figs. S7(e) and (j) for *P*_0_ = 25 and 32 respectively; *MSD* becomes slower as *T* decreases, typical of a glassy system. Note the qualitative difference of the cells at low-*P*_0_ and large-*P*_0_ at low *T*, the cell boundaries are compact in the former (Fig. S7(c)) whereas they become fractal-like in the latter (Fig. S7(h)).

**FIG S8.**
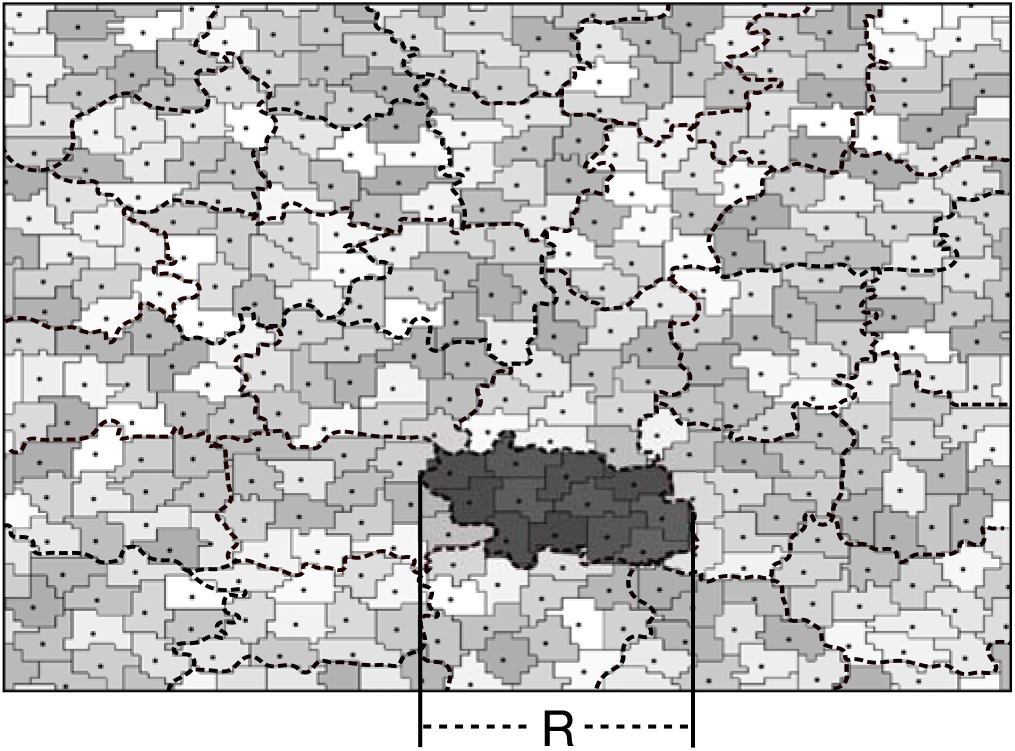
Schematic representation of the mosaic picture of RFOT theory. A glassy system consists of mosaics of different states as schematically shown by the dashed lines. Typical length scale, *ξ*, of the mosaics are given by two competing contributions, the configurational entropy and the surface reconfiguration energy.

### S3. EXTENDED RANDOM FIRST ORDER TRANSITION THEORY FOR CPM

As shown above as well as in the main text, the CPM has a disordered energy landscape. Therefore, we expect random first order transition (RFOT) theory phenomenology to be applicable for the glassy dynamics in such systems. Within RFOT theory, a glassy system consists of mosaics of different states as schematically shown in Fig. S8. Consider a region of length scale *R*, as shown by the shaded region in Fig. S8, in dimension *d* and look for the cost in energy for the rearrangement (changing state) of this region:

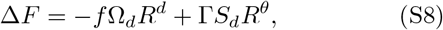

where *f* is the decrease in energy per unit volume due to the rearrangement, Ω_*d*_ and *S*_*d*_, volume and surface of a unit hypersphere, Γ, the surface energy cost per unit surface area due to the rearrangement and *θ* ≤ (*d* − 1) is the exponent relating surface area and length scale of a region. Within RFOT theory, the drive to reconfiguration is entropic in nature and given by the configurational entropy *s*_*c*_, that is *f* = *k*_*B*_*Ts*_*c*_, where *k*_*B*_ is the Boltzmann constant. Minimizing Eq. (S8), we get the typical length scale, *ξ*, for the mosaics as

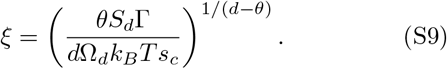

In general, the interaction potential, Φ, of the system determines *s*_*c*_ and Γ. In the case of CPM, the interaction potential is parameterized through *P*_0_, thus, Φ = Φ(*P*_0_). The temperature dependence of Γ is assumed to be linear [19], thus, Γ = Ξ[Φ(*P*_0_)]*T* and we write Eq. (S9) as

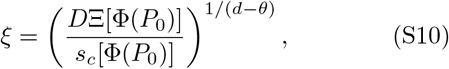

where *D* = *θS*_*d*_/*dk*_*B*_Ω_*d*_ is a constant. Within RFOT theory, relaxation dynamics of the system comes from relaxations of these individual mosaics of typical length scale *ξ*. The energy barrier associated for the relaxation of a region of length scale *ξ* is Δ(*ξ*) = Δ_0_*ξ*^*ψ*^, where Δ_0_ is an energy scale. The relaxation time then becomes *τ* = *τ*_0_ exp(Δ_0_*ξ*^*ψ*^/*k*_*B*_*T*), where *τ*_0_ is a microscopic time scale independent of *T*, but can depend on interatomic interaction potential, hence, on *P*_0_. Taking Δ_0_ = *κT*, where *κ* is a constant [20, 21] and setting *k*_*B*_ to unity, we obtain *τ* as

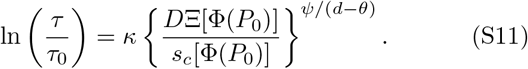

Following Refs. [21, 22] we take *θ* = *ψ* = *d*/2 and then Eq. (S11) can be written as

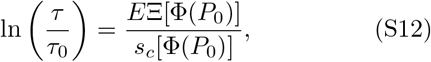

where *E* = *κD* is another constant. The theory presented here is similar in spirit with that for a network material obtained by Wang and Wolynes [23]. Our approach is perturbative in nature and we look at the effect of *P*_0_ by expanding the potential around a reference system. As discussed in the main text, the behavior of the system are different in the low-*P*_0_ and the large-*P*_0_ regimes that we discuss separately below.

#### Low-*P*_0_ regime

In this regime cells are not able to satisfy the perimeter constraint and the dynamics depends on *P*_0_. The simulation data for *P*_0_ = 24, …, 26.5 show sub-Arrhenius behavior and in the Angell plot representation, the curves tend towards the Arrhenius behavior as *P*_0_ decreases; the trend suggests existence of a particular value of *P*_0_ where the behavior becomes Arrhenius; we chose this 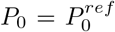 as the reference state and expand the effect of *P*_0_ on *s*_*c*_ and Ξ around this state. Thus, we have

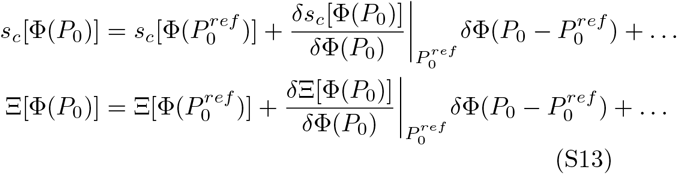

where we have ignored higher order terms. 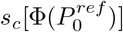, the configurational entropy for the reference system, vanishes at the Kauzmann temperature *T*_*K*_, thus,

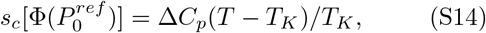

where Δ*C*_*p*_ is the difference of specific heats of the liquid and the periodic crystalline phase. It is evident from the discussion in Sec. S1 S1C [See Eq. (S7)], 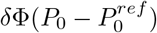, the change in potential due to a variation in *P*_0_, is proportional to 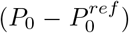. Our reference state is a confluent tightly packed state, any increase in *P*_0_ decreases repulsive interaction facilitating dynamics. This suggests higher *s*_*c*_ with increasing *P*_0_ while Ξ decreases. Therefore, we can write the first order terms as

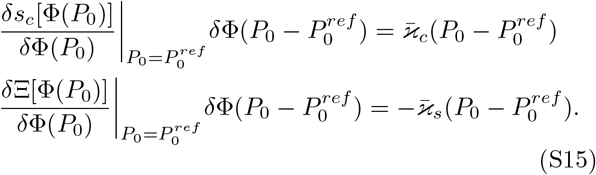

Using Eqs. (S13-S15) in Eq. (S12), we obtain

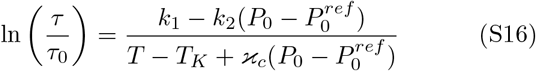

where 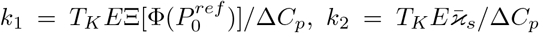 and 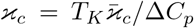 are all constants. The value of 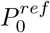 depends on the average cell area, for the results presented in the main text, the average cell area is 40 and we find 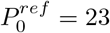 provides a good description of the data.

#### Large-*P*_0_ regime

The individual cells are able to satisfy the perimeter constraint in this regime, hence the actual value of *P*_0_ becomes irrelevant for the glassy dynamics and both *s*_*c*_ and Ξ become independent of *P*_0_. Therefore, we can simply write the RFOT theory expression for *τ* in this regime as

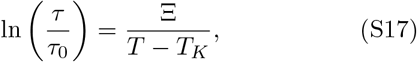

where the detailed values of Ξ and *T*_*K*_ for our system depend on whether *P*_0_ is even or odd since *P*_*i*_ can have even values only. Except this difference, the RFOT theory parameters, and hence the glassy dynamics, become independent of *P*_0_. However, the vestige of *P*_0_-dependence still remains in the dynamics through the high *T* properties of the system. We can not ignore the *P*_0_-dependence of *τ*_0_ in this regime, and, as we show in the main text [Fig. 3(d)], the difference in *τ* for different values of *P*_0_ can be understood in terms of *τ*_0_(*P*_0_).

#### System with *λ*_*P*_ = 0 and non-zero *J*

This system lacks any particular reference state and this makes the system different from that with *λ*_*P*_ ≠0. Perturbative expansion around a state with *J* = 0 is problematic and therefore, we need to chose a system with a moderate value of *J*. However, then any value of *J* is as good as any other. One consequence of this is that we expect the fragility of the system to be constant as discussed in the main text. Since the repulsive interaction only comes in the form of *J*, the trends of variation of *s*_*c*_ and Ξ with increasing *J* become opposite to that in the low-*P*_0_ regime above. Considering the reference system at a moderate value of *J*, we go through a similar set of arguments and obtain the RFOT expression for *τ* as

**FIG S9.**
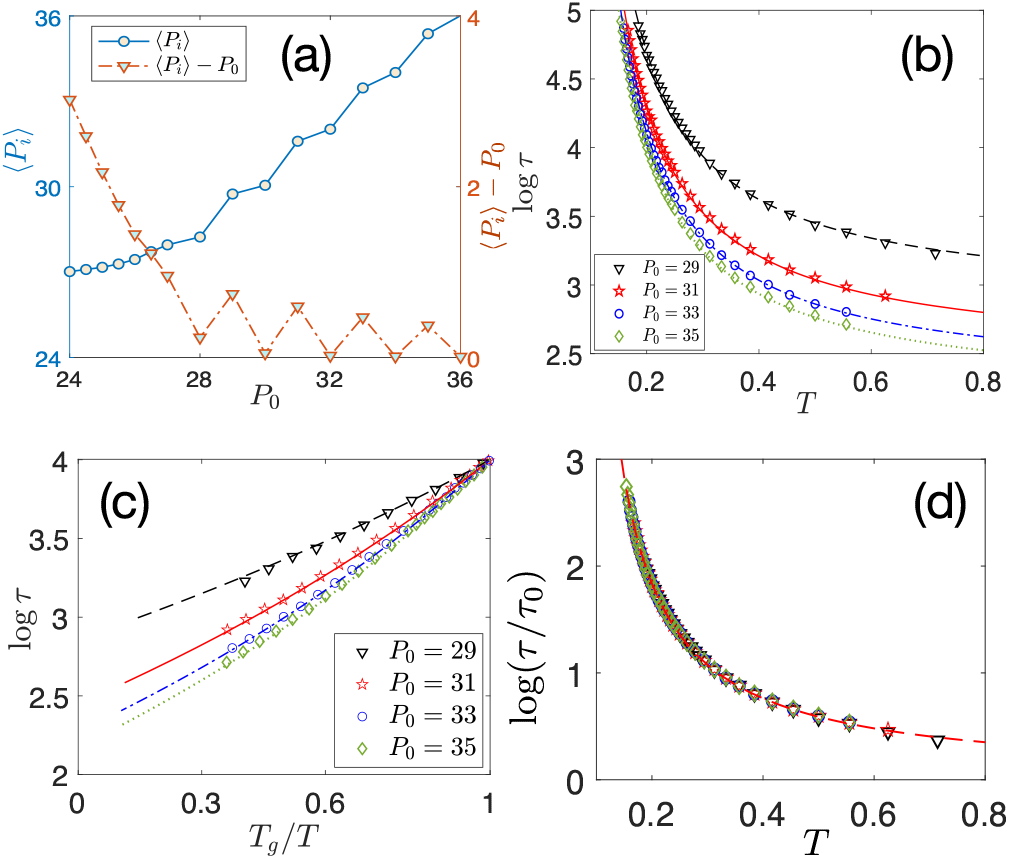
Behavior of CPM at the large-*P*_0_ regime for odd values of *P*_0_. (a) ⟨*P*_*i*_ ⟩ for odd values of *P*_0_ is slightly higher than that for even *P*_0_. Right *y*-axis shows ⟨*P*_*i*_ ⟩ − *P*_0_ as a function of *P*_0_ which goes to zero with increasing *P*_0_ with a slower rate than even *P*_0_. (b) Relaxation time *τ* as a function of *T* for odd values of *P*_0_. These data are well fitted with Eq. (6) in the main text with Ξ = 0.60 and *T*_*K*_ = 0.058 and *τ*_0_(*P*_0_) as a fitting parameter. *τ*_0_(*P*_0_) has a similar behavior (not shown) as for even values of *P*_0_ shown in the inset of Fig. (3b) in the main text. Lines are the RFOT theory fits. (c) *τ* in the Angell plot representation agrees well with the RFOT theory. (d) Plot of *τ*/*τ*_0_(*P*_0_) as a function of *T* for different values of *P*_0_ follows a master curve, implying that the glassiness for odd values of *P*_0_ is also independent of *P*_0_ in this regime.

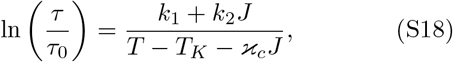

where *k*_1_, *k*_2_, *T*_*K*_ and ϰ_*c*_ are constants. Note the opposite signs of the constants *k*_2_ and ϰ_*c*_ in Eq. (S18) and Eq. (S16). Fitting Eq. (S18) with simulation data for *J* = 1, we obtain *k*_1_ = 0.284, *k*_2_ = 0.84, *T*_*K*_ = 0.04856 and ϰ_*c*_ = 0.157. Comparison of Eq. (S18) with the simulation data is presented in the main text, Figs. 4(c) and (d).

### S4. STRETCHING EXPONENT FOR THE DECAY OF *Q*(*t*)

It is well-known that the decay of self-overlap function *Q*(*t*) in a glassy system can be described through a stretched exponential function [24], the Kohlrausch-Williams-Watts (KWW) formula [25, 26] given by,

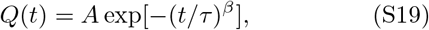

**FIG S10.**
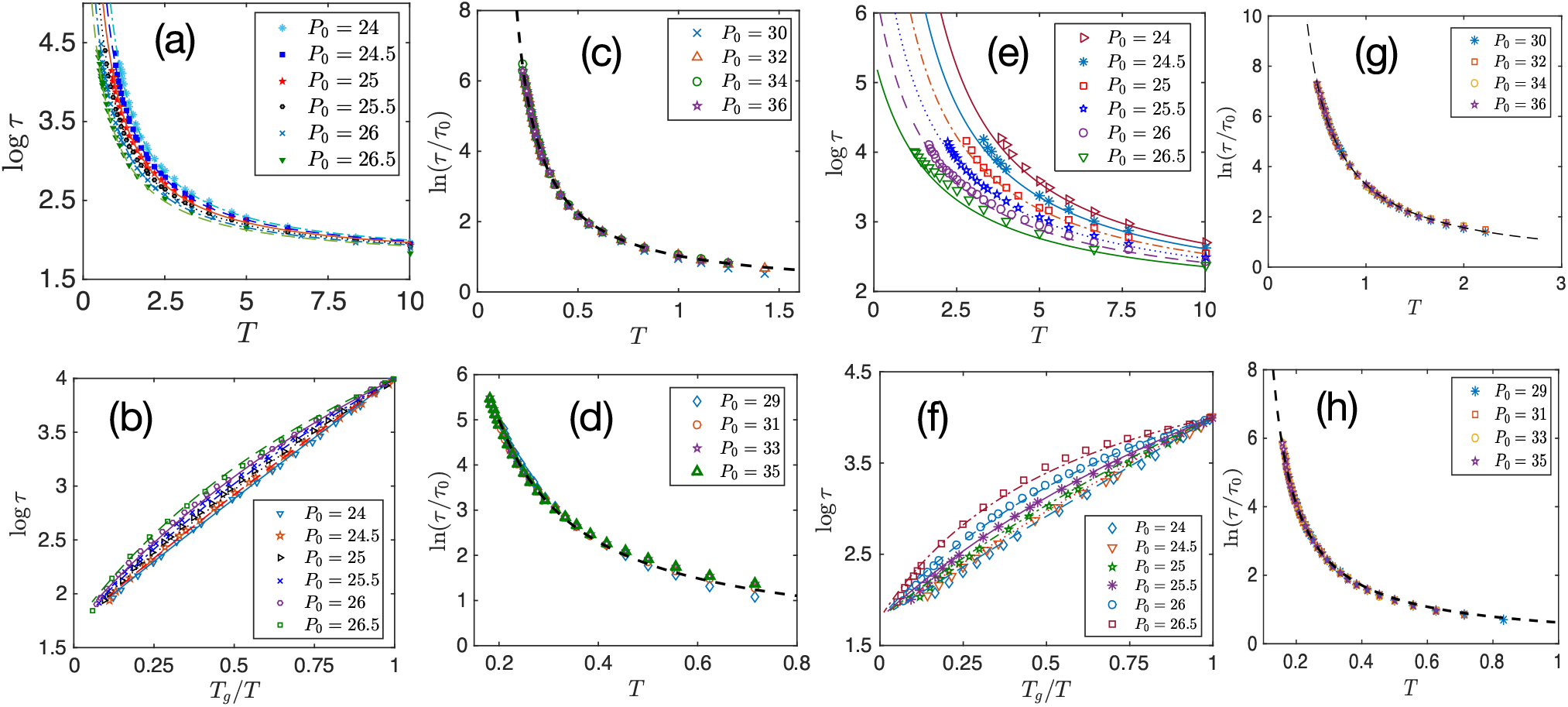
Results for *λ*_*P*_ = 0.25 (a-d) and *λ*_*P*_ = 1.0 (e-h). (a) *τ* as a function of *T* for different *P*_0_ with *λ*_*P*_ = 0.25. Symbols are simulation data and lines are RFOT theory, Eq. (S16), with the constants provided in Table S1. (b) Same data as in (a) but shown in the Angell plot representation. (c) *τ*/*τ*_0_ as a function of *T* for even-values of *P*_0_ in the large-*P*_0_ regime follow a master curve showing the *P*_0_-dependence in this regime comes from that in *τ*_0_. (d) Same as in (c) but for odd values of *P*_0_. (e-h) Corresponding plots as in (a-d) for *λ*_*P*_ = 1.0. Qualitative nature of the results shown here are similar with those for *λ*_*P*_ = 0.5 presented in the main text.

where *A* is a constant, of the order of unity, *τ*, the relaxation time and *β* is the stretching exponent. RFOT theory allows calculation of *β* through the fluctuation of local free energy barriers Δ*F* [27]. We assume that Δ*F* follows a Gaussian distribution given by,

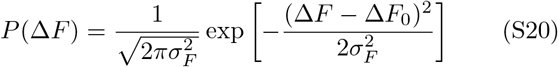

where Δ*F*_0_ is the mean of the distribution and *σ*_*F*_ is the standard deviation, which gives a measure of the fluctuation. Following Xia and Wolynes [27], we obtain *β* as

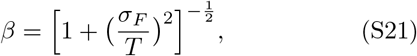

where we have set Boltzmann constant *k*_*B*_ to unity.

For the Gaussian distribution of Δ*F*, we obtain [27],

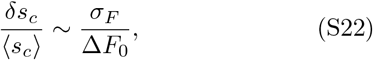

with 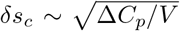, where *V* ∼ *ξ*^*d*^ is the typical volume of the mosaics. In the low-*P*_0_ regime, where we have compared our RFOT theory predictions with the simulation results, the length scale *ξ* of the mosaics, Eq. (S9), is given by,

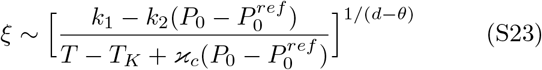

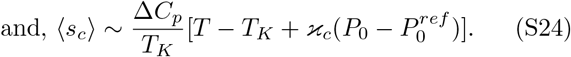

Using Eqs. (S23) and (S24), we obtain

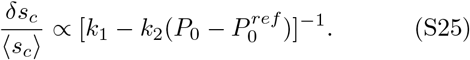

The mean free energy barrier (Δ*F*_0_) is obtained, by using *R* = *ξ* in Eq. (S8), as

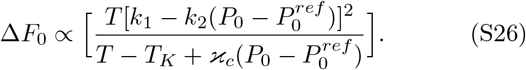

Using Eqs. (S25), (S26) and (S22) in Eq. (S21), we obtain *β* as

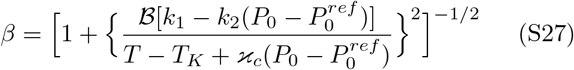

Where ℬ is a constant. It is well-known that RFOT theory predicts the correct trends of *β*, but the absolute values differ by a constant factor even for a particulate system [27]. Since we are interested in the trend of *β* as a function of *P*_0_, we multiply Eq. (S27) by a constant 𝒜 to account for this discrepancy and obtain

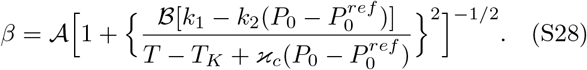

The constants *k*_1_, *k*_2_, *T*_*K*_ and ϰ_*c*_ are already determined, 𝒜 and ℬ are obtained through the fit of Eq. (S28) with the simulation data for *P*_0_ = 25 as a function of *T*.

The decay of *Q*(*t*) at high *T* is stretched exponential followed by a power law at long times. To avoid this high *T* power-law regime in the analysis, we fit the data up to a time *τ*, that is when *Q*(*t*) = 0.3 with Eq. (S19) and obtain *β* as shown in Figs. 4(e) and (f) in the main text.

### S5. DYNAMICS IN THE LARGE-*P*_0_ REGIME FOR ODD VALUES OF *P*_0_

For simplicity of presentation, we showed the results in the large-*P*_0_ regime for even values of *P*_0_ alone. We now present the results for odd values of *P*_0_ in this regime. As stated in Sec. S2, the qualitative behaviors for odd *P*_0_ are similar to those with even *P*_0_, however, since the cell perimeter can only be even due to the underlying lattice, ⟨*P*_*i*_⟩ − *P*_0_ goes to zero with increasing *P*_0_ at a slower rate than that for even *P*_0_ as shown in Fig. S9(a). The data for relaxation time *τ* as a function of *T* is shown in Fig. S9(b) by symbols and the corresponding RFOT theory fits of Eq. (6) in the main text are shown by lines with Ξ = 0.60 and *T*_*K*_ = 0.058; *τ*_0_(*P*_0_) has a similar behavior as shown in the inset of Fig. (3b) in the main text. The Angell plot representation, as shown in Fig. S9(c) of the same data as presented in Fig. S9(b), agrees well with the RFOT theory. Finally, to show that glassiness in this regime for odd values of *P*_0_ is also independent of *P*_0_, we plot *τ*/*τ*_0_ as a function of *T* and find excellent data collapse for different values of *P*_0_ as shown in Fig. S9(d) and the master curve agrees well with the RFOT theory (line).

### S6. DYNAMICS FOR DIFFERENT *λ*_*P*_

The qualitative behavior of the system for different values of *λ*_*P*_ remains same, although the quantitative values of the parameters within the RFOT theory description depend on *λ*_*P*_. For the results presented in the main text, we have used *λ*_*P*_ = 0.5; here we present the results for *λ*_*P*_ = 0.25 in Figs. S10 (a-d) and for *λ*_*P*_ = 1.0 in Figs. S10 (e-h). For the two values of *λ*_*P*_, we show *τ* as a function of *T* in the low-*P*_0_ regime for different *P*_0_ in Figs. S10(a) and (e) and the corresponding Angell plots in Figs. S10(b) and (f); the unusual sub-Arrhenius behavior, found for *λ*_*P*_ = 0.5 as shown in the main text (Fig. 2e) is also present at these two different values of *λ*_*P*_. The glassy dynamics becomes independent of *P*_0_ in the large-*P*_0_ regime as confirmed by the data collapse for *τ*/*τ*_0_ as a function of *T*, shown in Figs. S10(c) and (d) for even and odd values of *P*_0_ respectively for *λ*_*P*_ = 0.25 and in Figs. S10(g) and (h) for even and odd values of *P*_0_ respectively for *λ*_*P*_ = 1.0. This implies any *P*_0_-dependence in this regime must come from that of *τ*_0_(*P*_0_). The symbols in Fig. S10 are simulation data and the lines are RFOT theory predictions, Eq. (S16), with the constants given in Table S1.

**FIG S11.**
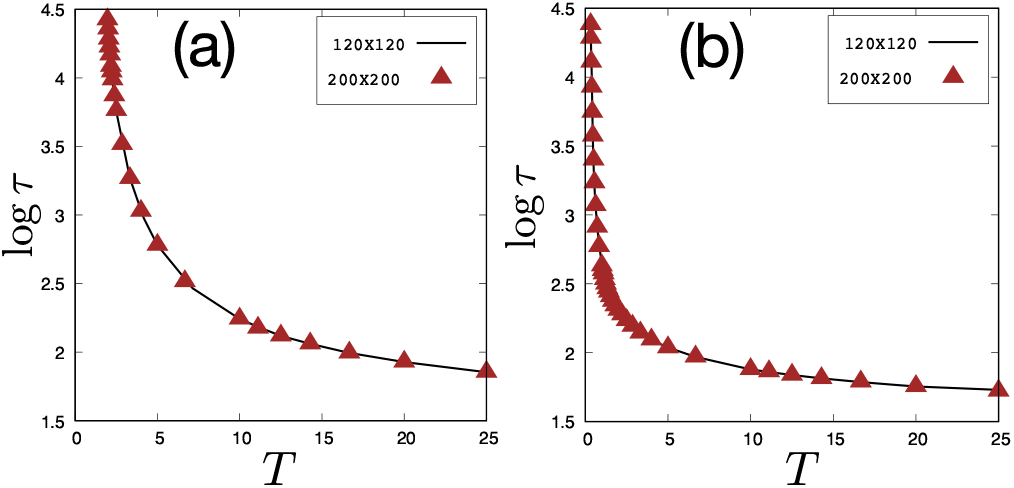
Finite system size effects are negligible in our results. We plot *τ* as a function of *T* for two different systems of sizes 120 × 120 and 200 × 200 for *P*_0_ = 24 in (a) and *P*_0_ = 32 in (b). Data for the two systems are essentially identical. Cell sizes of average area 40 are same in these two systems.

**TABLE S1.** Since the average cell size remains same in all the systems, we use 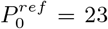. Values of the constants *k*_1_, *k*_2_, ϰ_*c*_ and *T*_*K*_, appearing in Eq. (S16), for different values of *λ*_*P*_ are given below.

| $\lambda_P$ | $k_1$ | $k_2$ | $\varkappa_c$ | $T_K$ |
| --- | --- | --- | --- | --- |
| 0.25 | 7.44627 | 0.610723 | 0.146849 | 0.079553 |
| 0.5 | 14.7834 | 1.21128 | 0.307531 | 0.005677 |
| 1.0 | 22.9153 | 2.21207 | 0.685 | 0.5828 |

### S7. EFFECT OF FINITE SYSTEM SIZES ON THE DYNAMICS

To investigate the effect of finite sizes of the system on the dynamics, we have looked at several systems of different sizes *L* × *L* where *L* varies from 80 to 200 keeping the sizes of the cells same, i.e., of area 40. We find that the results for the system size 120 × 120 that we have mainly investigated and presented the results in the main text remain unchanged when we use larger system sizes. For a comparison, we show *τ* as a function of *T* for two different systems in Fig. S11 for *P*_0_ = 24 and 32, the data for the two systems are essentially same.

### S8. EFFECT OF DIFFERENT CELL SIZES

We have shown in the main text that the two different regimes result due to geometric constraint where the cells can not fully satisfy the perimeter constraint in the low-*P*_0_ regime whereas they are able to satisfy it in the large-*P*_0_ regime. As we vary the average size of the cells in the system, numerical value of *P*_0_ for the transition from one regime to the other changes, but apart from that, the qualitative behavior of the system remains same. We have verified this with different systems of various cell sizes. We present simulation data of *τ* as a function of *T* for a system of size 180 × 180 with average cell area 90 and 360 total number of cells in Fig. S12 (a) and the same data in Angell plot representation in Fig. S12 (b). We have used *λ*_*P*_ = 0.5 for these simulations to compare the results with those presented in the main text. For average cell area 90, the minimum possible perimeter is 38, thus, we expect the geometric transition point, separating the low-*P*_0_ regime and the large-*P*_0_ regime, to be somewhere between 39 and 40. Lines in Fig. S12 represent plots of RFOT theory, Eq. S16, with the parameters appearing in Eq. (S16) as follows: 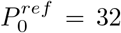, *τ*_0_ = 68.72, *k*_1_ = 27.46, *k*_2_ = 1.39, ϰ_*c*_ = 0.29 and *T*_*K*_ = 0.073. We have checked (data not presented) that glassiness in the large-*P*_0_ regime becomes independent of *P*_0_. Thus, the main features of glassiness in this system are similar to those for the system with average cell area 40 as presented in the main text.

**FIG S12.**
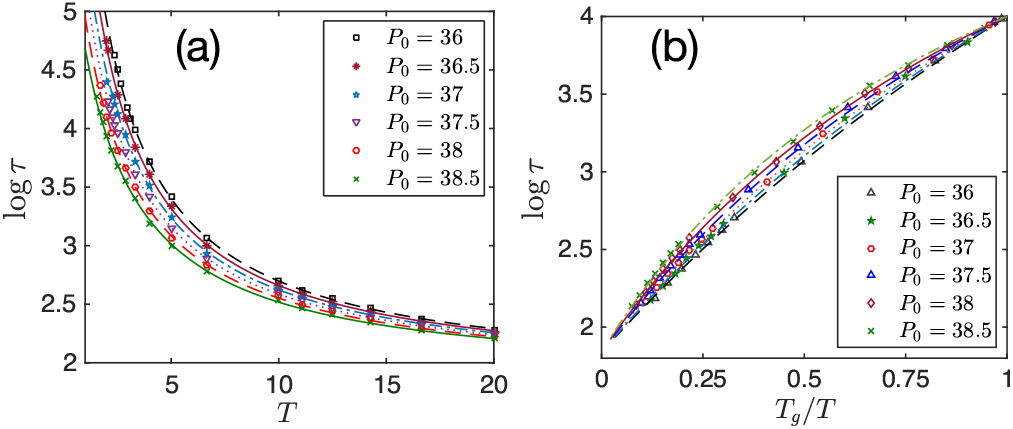
Systems with varying cell sizes have different average cell areas. This sets the value of *P*_0_ separating the low-*P*_0_ and large-*P*_0_ regimes, as it is a geometric effect. Except this, the qualitative behavior of the system remains same for different cell sizes. (a) *τ* as a function of *T* for different *P*_0_, (b) same data as in (a) in the Angell plot representation. Symbols are simulation data and lines are the RFOT theory, Eq. (S16), with the constants quoted in the text.

**FIG S13.**
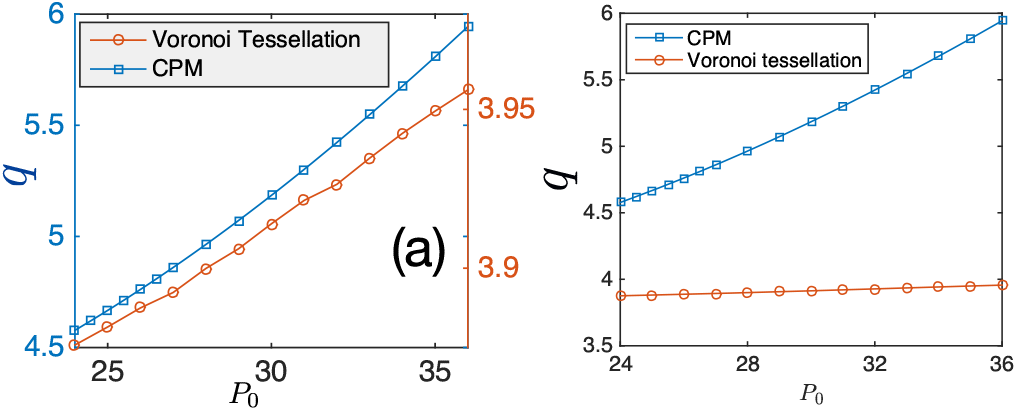
Observed shape index, 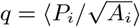, for the CPM and that obtained via the voronoi tessellation of the centers of mass of the cells. (a) Two estimates are plotted in two different scales to emphasize their qualitatively similar behaviors as a function of *P*_0_. (b) They are plotted on the same scale to highlight that voronoi tessellation underestimates the value of *q* in comparison to its actual value in CPM, the discrepancy becomes larger at higher *P*_0_.

### S9. DIFFERENCE OF CPM AND VERTEX-BASED MODELS

There are different ways of representing the value of the perimeter of different cells in CPM. One of the methods, often opted, is to coarse-grain over a number of sites and represent the boundary as a smooth curve [2]. As coarse-graining obscures some of the microscopic features of the model, we have presented the perimeter as the actual boundary. On a square lattice, the minimum perimeter configuration for a certain area is a square, thus the quantitative values of the parameters will differ between CPM and its continuum versions, the vertex-based models, though they all represent a confluent system. For a comparison, we show the observed shape index 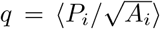 for different values of *P*_0_ at a fixed *T* = 5.0 in Fig. S13 obtained from two different estimates as follows: first, we obtain the actual values of *q* in the CPM, second, we construct a voronoi tessellation for the centers of mass of the cells and obtain *q* for that system. Although the qualitative trends for both estimates are similar, that is, they both monotonically increase with *P*_0_ as shown in Fig. S13(a) (note the difference in scales), the voronoi tessellation underestimates *q* as shown in Fig. S13(b); the discrepancy becomes higher at larger *P*_0_ where the voronoi tessellation can not account for nonlinear cell boundaries.

## Footnotes

1 This statement is strictly true for a continuous system. In our lattice model though, there is an upper limit of 2*A*_*i*_ +2, where *A*_*i*_ is the area. Since this upper limit value is very large compared to the regime of our investigation, we expect this upper limit to be irrelevant.

